# Unifying the analysis of bottom-up proteomics data with CHIMERYS

**DOI:** 10.1101/2024.05.27.596040

**Authors:** Martin Frejno, Michelle T. Berger, Johanna Tüshaus, Alexander Hogrebe, Florian Seefried, Michael Graber, Patroklos Samaras, Samia Ben Fredj, Vishal Sukumar, Layla Eljagh, Igor Brohnshtein, Lizi Mamisashvili, Markus Schneider, Siegfried Gessulat, Tobias Schmidt, Bernhard Kuster, Daniel P. Zolg, Mathias Wilhelm

**Author notes:** **Corresponding authors:** Martin Frejno, Mathias Wilhelm. Contributed equally (Martin Frejno, Michelle T. Berger, Johanna Tüshaus).

## Abstract

Proteomic workflows generate vastly complex peptide mixtures that are analyzed by liquid chromatography-tandem mass spectrometry (LC-MS/MS), creating thousands of spectra, most of which are chimeric and contain fragment ions from more than one peptide. Because of differences in data acquisition strategies such as data-dependent (DDA), data-independent (DIA) or parallel reaction monitoring (PRM), separate software packages employing different analysis concepts are used for peptide identification and quantification, even though the underlying information is principally the same. Here, we introduce CHIMERYS, a novel, spectrum-centric search algorithm designed for the deconvolution of chimeric spectra that unifies proteomic data analysis. Using accurate predictions of peptide retention time, fragment ion intensities and applying regularized linear regression, it explains as much fragment ion intensity as possible with as few peptides as possible. Together with rigorous false discovery rate control, CHIMERYS accurately identifies and quantifies multiple peptides per tandem mass spectrum in DDA, DIA and PRM experiments.

## Introduction

Mass spectrometry-based bottom-up proteomics is the mainstay technology for high-throughput protein identification and quantification today^1–3^. The former is achieved by matching theoretical, predicted or library fragment ion mass spectra (MS2) to experimental MS2 spectra, which contain sequence and amino acid modification information on peptide precursor ions, measured in MS1 spectra. Today, MS2 spectra are typically acquired in data-dependent (DDA), data-independent (DIA) or parallel reaction monitoring (PRM) mode. Peptide quantification either uses the peptide ion intensity from MS1 (DDA) or fragment ion intensities from MS2 (DIA, PRM) spectra. A central challenge for data analysis lies in the fact that most MS2 spectra are chimeric, i. e. they contain more than one peptide because LC-MS/MS systems cannot fully separate the vast number of peptides resulting from whole proteome enzymatic digestion.

DIA MS2 spectra are usually more complex than DDA MS2 spectra because they are typically acquired with wider isolation windows to maintain low MS cycle times (important for quantification) and hence contain fragment ions from many different precursors^4^. Although DDA and PRM MS2 spectra are typically acquired to minimize co-isolation, they are also chimeric, albeit to a much lesser extent^5^. Because of the way different data acquisition approaches have evolved, the resulting data types are analyzed differently^6^, making it difficult to compare them in an unbiased fashion^7^.

DDA data is analyzed in a so-called spectrum-centric fashion^6^. Database search algorithms for DDA data attempt to maximize identifications from chimeric spectra by submitting them multiple times using several precursors detected in the isolation window. Often, fragment ions explained by a given peptide are removed from the spectrum before it is searched again^5,8^. While often able to call a second or third peptide, this approach will lead to an underutilization of the spectral information when fragments are shared between peptides, resulting in reduced sensitivity. In case fragment ions are not removed for an additional search, there is a danger that the same information is used too often, resulting in reduced specificity. In any case, the central output of DDA search engines is one or multiple peptide-spectrum matches (PSMs) per experimental MS2 spectrum.

In contrast, DIA and PRM data analysis follows a so-called peptide-centric approach that asks the question if any of a pre-defined list of peptides are detectable in extracted ion chromatograms (XICs) of their MS1 and/or MS2 spectra^6,9^. This approach requires the use of peptide spectral libraries, which can be generated from previous experimental data, predicted via machine or deep learning models, constructed directly from the DIA data itself by scoring (deconstructed) MS2 spectra in a DDA-like fashion^10^ or a combination of these approaches. Subsequently, the queried peptides are detected and quantified by extracting co-eluting fragment ion chromatograms based on the spectral library.

Because of the molecular complexity of proteomic samples and the large quantities of MS2 spectra of varying quality that are generated by LC-MS/MS, accurate false discovery rate (FDR) control is an important part of data analysis, particularly in large-scale projects. While FDR control for DDA data is rather mature^11–14^, it is still a substantial challenge in DIA data, because constructing realistic decoy MS2 spectra and retention times is far from obvious, an issue increasingly realized and addressed by machine learning algorithms for peptide property prediction^15–17^.

In this work, we introduce a novel spectrum-centric and data acquisition-agnostic approach for the analysis of MS2 spectra, implemented in the search algorithm CHIMERYS. It deconvolutes any MS2 spectrum, regardless of whether it was acquired by DDA, DIA or PRM, thus unifying the analysis of bottom-up proteomics data. We build upon a concept introduced for the deconvolution of DIA spectra using spectral libraries^4^ and leverage deep learning-based predictions of fragment ion intensities in conjunction with linear algebra for the deconvolution of MS2 spectra. The resulting signal contributions of each peptide identified in each MS2 spectrum can be combined into a quantitative readout. Applying the approach substantially enhances identification rates of PSMs, peptides, and proteins across all sample types in DDA, enables the hands-off processing of PRM data and matches the performance of alternative DIA software while maintaining accurate FDR control throughout.

## Results

### Deconvolution of chimeric DDA spectra

The core assumption behind CHIMERYS is that chimeric MS2 spectra are linear combinations of pure spectra from co-isolated precursors. The algorithm is entirely spectrum-centric and employs non-negative L1-regularized regression via the LASSO^18^ to explain as much experimental intensity as possible with as few peptide precursors as possible (Figure 1A). It uses highly accurate predictions of fragment ion intensities and retention times for target and decoy peptides instead of spectral libraries.

**Figure 1.**
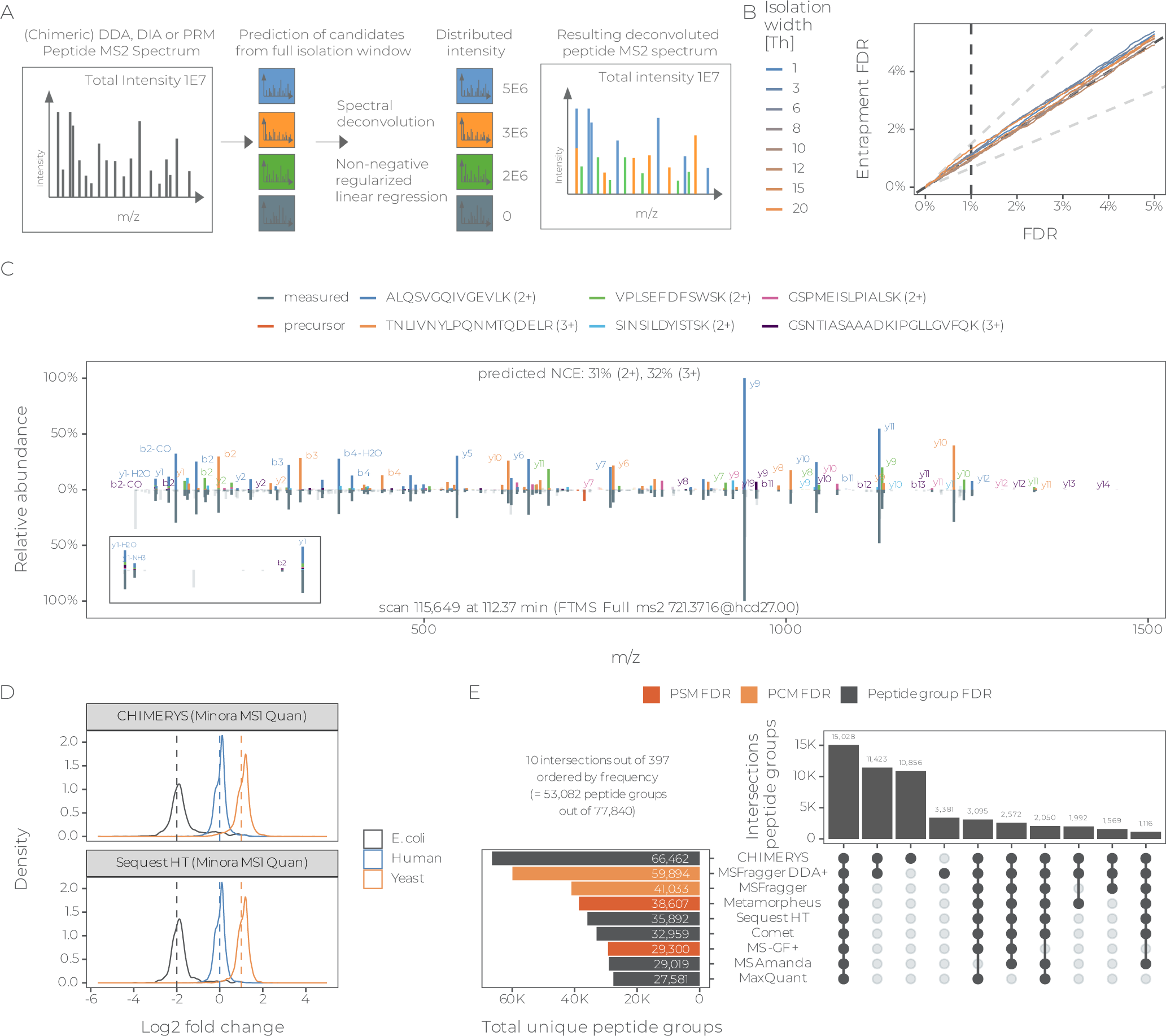
Deconvolution of chimeric DDA spectra: **(A)** CHIMERYS treats chimeric spectra as linear combinations of pure spectra, and its spectrum-centric deconvolution uses non-negative regularized regression to estimate interference-corrected total ion currents for all candidate peptides. **(B)** Peptide-level entrapment analysis with the classic eFDR approach (see Online Methods) of DDA data acquired using different isolation windows widths and processed with CHIMERYS. **(C)** Example of a deconvoluted chimeric spectrum with six PSMs from a 2-hour HeLa DDA single-shot measurement, acquired on an Orbitrap QE HF-X with 1.3 Th isolation windows from the LFQbench-type dataset^22^. The inset visualizes how the experimental intensity of shared fragment ions with low m/z values is distributed to multiple PSMs. **(D)** Peptide-level density plots for triplicate 2-hour DDA single-shot measurement from two different conditions, acquired on an Orbitrap QE HF-X with 1.3 Th isolation windows from the LFQbench-type dataset, analyzed with CHIMERYS (top) or Sequest HT (bottom). FDR was controlled at 1% at the global peptide group-level. **(E)** Comparison of peptide identifications from multiple search engines on the same data as in **(C)**. FDR was natively controlled at different levels, depending on the search engine.

Briefly, predicted MS2 spectra from precursors with predicted retention times that fall within a data-dependent retention time window and precursor isotope envelopes that (partially) overlap with the isolation window are compared to experimental MS2 spectra based on a multitude of fragment ion intensity-free and -dependent scores for each PSM (Online Methods). Next, spurious PSMs are removed based on some of these scores. For example, PSMs are required to have at least three matched fragment ions, one of which must be the base peak (most abundant peak of the prediction) and another one of which must be among the top three most intense peaks of the predicted spectrum. PSMs passing these criteria are used for deconvolution, where they compete for experimental fragment ion intensity in one concerted step; an approach fundamentally different from the classic subtraction methods (Figure 1A). PSMs with enough contribution to the experimental spectrum as measured by CHIMERYS coefficients and that pass additional score filters are handed to mokapot^13^ for PSM-level FDR control, specifically allowing for multiple PSMs per spectrum, similar to DIAmeter^19^.

We validated this FDR estimation on data with varying chimericity by systematically increasing the isolation window widths of 1-hour HeLa single-shot measurements from 1.4 up to 20.4 Th using entrapment experiments (Online Methods). Figure 1B shows that CHIMERYS’ peptide-level q-values correspond to empirical q-values calculated based on entrapment identifications with the classic eFDR approach, independent of isolation window width.

Figure 1C displays the confident identification of six peptides with relative contributions to the experimental total ion current ranging from 4% to 54% in a mirror spectrum. Notably, the experimental intensities for the y1, y1-NH_3_ and y1-H_2_O ions that are shared between the five peptides (C-terminal lysine) align well with the sum of predicted intensities of the corresponding peptides, scaled by their respective CHIMERYS coefficient, which can be interpreted as the interference-corrected total ion current of a peptide in an MS2 spectrum (Online Methods). This exemplifies how the algorithm identifies multiple peptides in chimeric spectra while distributing intensities of shared fragment ions. Peptides identified by CHIMERYS recapitulate the expected quantitative ratios in a multi-organism-mixture experiment (Figure 1D). This renders CHIMERYS suitable for approaches like wide-window DDA acquisition (also termed WWA or wwDDA)^20,21^ and the direct analysis of DIA data.

To assess the performance of the algorithm on DDA data, we analyzed a 2-hour HeLa cell lysate digest with 1.3 Th MS2 isolation windows (Online Methods). CHIMERYS identified 238,795 PSMs at 1% PSM FDR with an overall identification rate of >85% (Supplementary Figure 1A). More than two thirds of identified MS2 spectra contained more than one precursor (Supplementary Figure 1B) confirming previous observations^5^. Fragment ions shared between different peptides were detected across the full MS2 m/z range with an expected higher frequency <200 m/z (Supplementary Figure 2), rendering current approaches for handling chimeric spectra error prone. Comparing these results to seven academic and commercial DDA search engines (Figure 1E) revealed that CHIMERYS identifies many additional peptides (Supplementary Figure 3). Most of these additional identifications stem from low abundant peptides (Supplementary Figure 4A) with fewer matched fragment ions (Supplementary Figure 4B) that were identified based on intensity-dependent scores such as the normalized spectral contrast angle (Supplementary Figure 4C). This resulted in a markedly higher number of peptides per protein group in CHIMERYS compared to Sequest HT (Supplementary Figure 4D). It is worth noting that some of these search engines do not control FDR at the same level, which has a substantial influence on such comparisons (Supplementary Figure 4E-F). Controlling FDR at a ‘lower’ level and counting identifications at a ‘higher’ level (e.g. counting peptides at PSM FDR) will overestimate the number of identifications. Identifications need to be reported at the same level at which FDR is controlled.

The gains observed for HeLa digests relative to Sequest HT were corroborated using more difficult biological samples at the protein group level (urine: +21%; CSF: +17%; plasma: +10%; FFPE material: +37%; secretomes: between +33% and +71%, *Arabidopsis thaliana*: +13%; *Haliobacterium*: +20%) (Supplementary Figure 5A-F). This data highlights that CHIMERYS substantially increases the analysis depth of DDA data without changing data acquisition.

### Revisiting legacy data using CHIMERYS

We conducted a retrospective study of HeLa single-shot analyses spanning many years and Orbitrap instrument generations. Despite many differences that impair a truly fair comparison, a clear trend was observed in that the higher the speed and sensitivity of the instrument, the higher the advantage of CHIMERYS over Sequest HT (Figure 2A, Supplementary Figure 6A).

**Figure 2.**
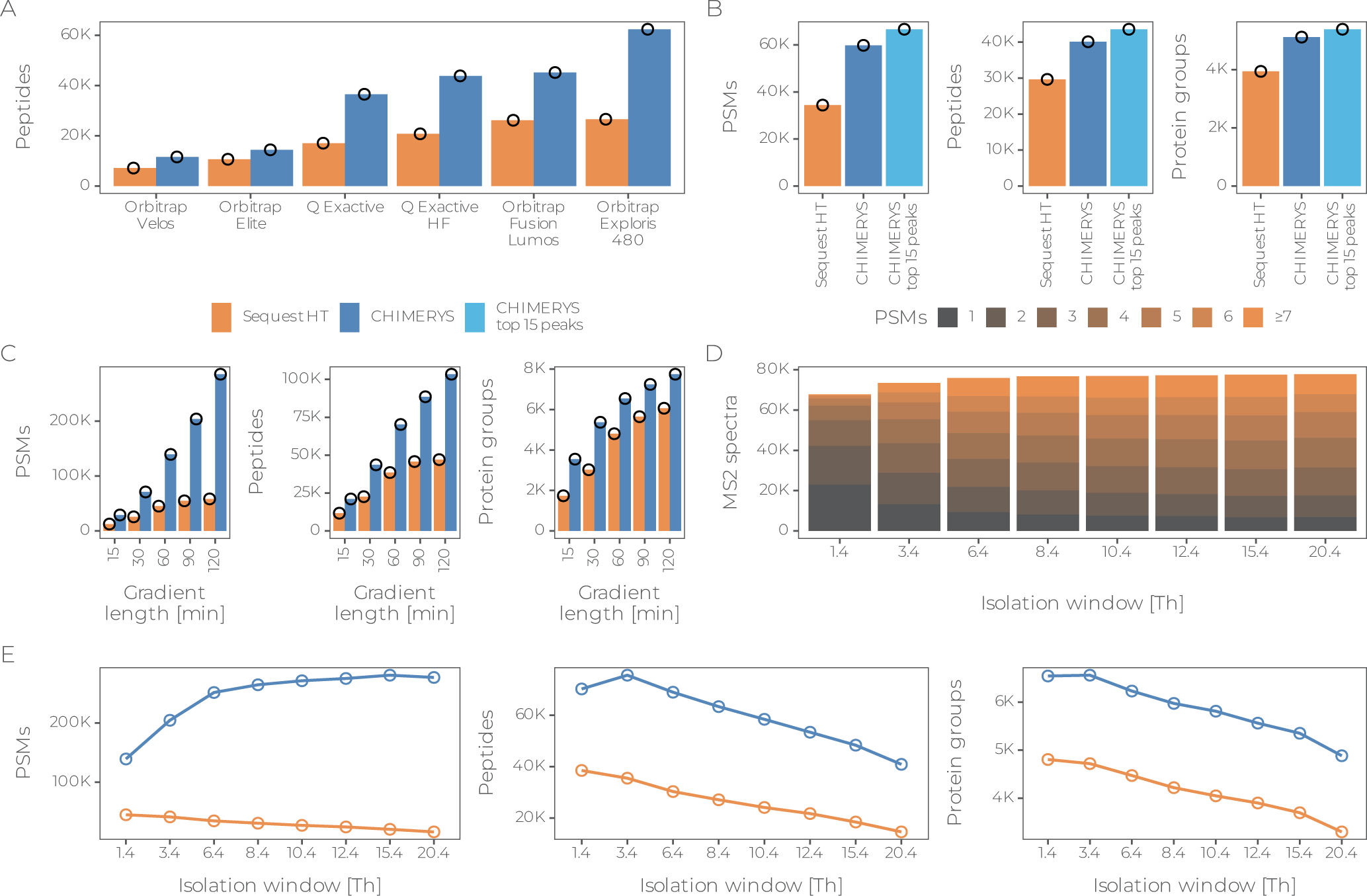
Optimizing data acquisition with deconvolution in mind. **(A)** Peptide identifications at 1% global peptide group-level FDR based on Sequest HT (orange) and CHIMERYS (blue) from 1h HeLa single-shot measurements, acquired using various Orbitrap generations. **(B)** PSM, peptide and protein group identifications based on Sequest HT (orange), CHIMERYS (dark blue) and CHIMERYS after removal of low-abundance peaks (light blue) from a 1-hour HeLa single-shot measurement, acquired using CID fragmentation with ion trap read out. FDR was controlled at 1% at the run-specific PSM-, global peptide group- and global protein level, respectively. **(C)** PSM, peptide and protein group identifications based on Sequest HT (orange) and CHIMERYS (blue) from HeLa single-shot measurements using various gradient lengths. FDR was controlled at 1% at the run-specific PSM-, global peptide group- and global protein level, respectively. **(D)** Distribution of the number of PSMs per MS2 spectrum from 1-hour HeLa single-shot measurements, acquired using different isolation window widths. FDR was controlled at 1% at the run-specific PSM-level. **(E)** PSM, peptide and protein group identifications based on Sequest HT (orange) and CHIMERYS (blue) from 1-hour HeLa single-shot measurements, acquired using different isolation window widths. FDR was controlled at 1% at the run-specific PSM-, global peptide group- and global protein level, respectively.

Next, we investigated low-resolution ion trap data (ITMS) comparing CHIMERYS to Sequest HT on unprocessed spectra and on spectra filtered for containing only the top 15 most abundant fragments per 100 Th window (Figure 2B). In contrast to high-resolution Orbitrap data, we observed a notable improvement by removing low-abundance peaks in ITMS spectra. Specifically, CHIMERYS identified 74% more PSMs, 35% more peptides, and 30% more protein groups compared to Sequest HT on unprocessed spectra were, while it identified 94% more PSMs, 47% more peptides, and 37% more protein groups on spectra preprocessed with a top 15 by 100 Th filter. Both examples show that substantially more information can be extracted from legacy data by harnessing the information contained in chimeric spectra.

### Optimizing data acquisition with deconvolution in mind

We next assessed to what extent CHIMERYS’ capability to deconvolute highly complex spectra can be used to optimize data acquisition. First, we evaluated LC gradients with the goal to increase sample throughput per day (SPD; Online Methods). Figure 2C shows that CHIMERYS identified a similar number of peptides and proteins in 30 min (48 SPD) for which Sequest HT needed 120 min (12 SPD) of the same HeLa digest, increasing throughput by a factor of four.

Next, we explored a possible increase in identification efficiency by intentionally widening the isolation windows in DDA (between 1.4 Th to 20.4 Th; Supplementary Figure 6B-D). The analysis revealed that the number of identified PSMs increased with wider isolation windows (Figure 2E) and began to plateau at > 8 m/z. This is likely due to the AGC limit, which – together with the dynamic range of MS2 spectra – limits the number of peptides in chimeric spectra with a sufficient number of detectable fragment ions. The number of peptide (and protein) identifications reached its maximum at a window size of 3.4 Th for this specific dataset and substantially decreased for larger isolation windows, likely due to the fact that more and more of the limited number of PSMs were from the same, high-abundant peptides. Such approaches have also gained popularity in single-cell proteomics (SCP), where extended injection times enhance sensitivity but result in fewer MS2 scans. CHIMERYS can counteract this effect and was already applied to extract more PSMs and peptides from these intentionally chimeric scans in SCP data^21^. The strong gains at the PSM, peptide and protein level are noteworthy as identifying nearly 8,000 proteins in a single 120 min DDA analysis has rarely been achieved before and implies that chromatographic pre-fractionation of samples may no longer be necessary.

### Deconvolution of chimeric DIA spectra

CHIMERYS deconvolutes DIA spectra in the same way as described for DDA spectra above. The only difference is that DIA spectra are usually more chimeric. Exemplified by a high-load LFQbench-type multi-organism mixture dataset^22^, CHIMERYS identified an average of 529,993 PSMs per raw file at 1% run-specific PSM FDR, mapping to 66,888 unique peptide groups and 7,331 unique protein groups at 1% global peptide group and protein FDR, respectively, with an overall identification rate of >60% (Supplementary Figure 7A). More than 82% of identified MS2 spectra contained more than one precursor (Supplementary Figure 7B) and shared fragment ions were more frequent, emphasizing the need for spectrum deconvolution that assigns shared fragment ions *pro rata* to the contributing peptides (Supplementary Figure 7C-J).

### Comparison to other DIA search engines

We compared results obtained with CHIMERYS on DIA data acquired on an Orbitrap QE HF-X from the LFQbench-type dataset to the library-free workflows implemented in the popular software tools DIA-NN^23^ and Spectronaut^24^ using entrapment experiments to validate FDR control in the run-specific context^25^ (see Online Methods for context definitions and search parameters). The results show that CHIMERYS’ self-reported q-values correspond to the empirical q-values calculated based on entrapment identifications (Supplementary Figure 8A). DIA-NN and Spectronaut appeared to underestimate FDR based on all three or the peptide and concatenated entrapment approaches, respectively (Supplementary Figure 8B-C). Recently proposed more stringent settings for Spectronaut^26^ had little if any effect on this issue (Supplementary Figure 8D). Similar observations were made when analyzing the TimsTOF Pro data of the LFQbench-type dataset using Spectronaut (Supplementary Figure 8E). All analyses below used the peptide eFDR approach (eFDR from here onwards). Filtering on eFDR in addition to the algorithm-dependent self-reported FDR did not change the overall number of identifications for CHIMERYS. Reductions to a level comparable to CHIMERYS’ results were observed for DIA-NN and Spectronaut. Data completeness for CHIMERYS did not change when requiring precursors to be quantified in two out of three replicate experiments. Reductions to a level similar to CHIMERYS’ results were observed for Spectronaut and to a level below CHIMERYS for DIA-NN (Figure 3A). At full data completeness, CHIMERYS and Spectronaut substantially outperformed DIA-NN on the LFQbench-type data (Supplementary Figure 9A).

**Figure 3.**
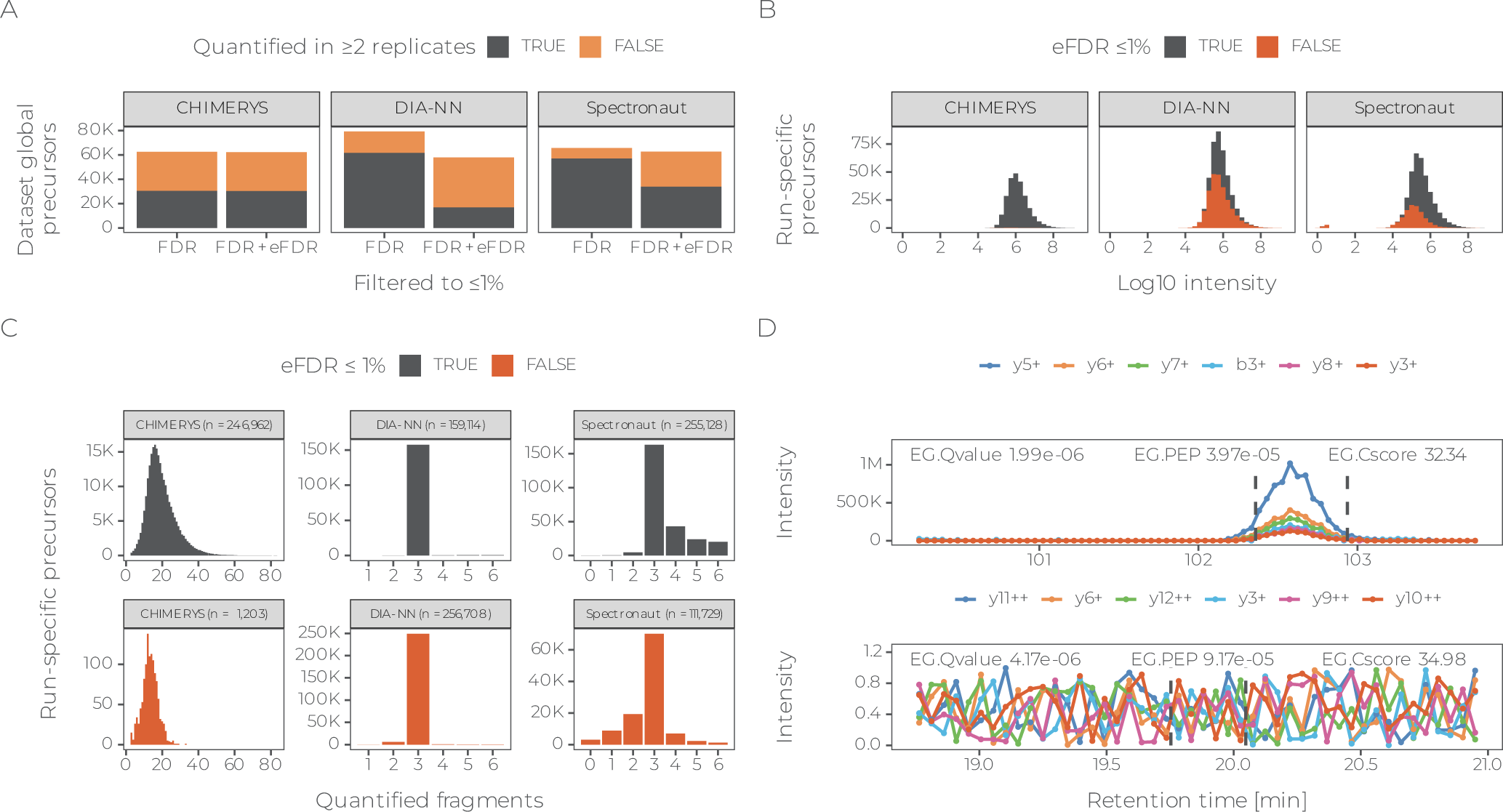
Deconvolution of chimeric DIA spectra. **(A)** Precursors quantified by CHIMERYS, DIA-NN and Spectronaut in at least one (orange) or two (gray) out of three replicate measurements of two different conditions in triplicate 2-hour DIA single-shot measurement from two different conditions, acquired on an Orbitrap QE HF-X with 8 Th isolation windows from the LFQbench-type dataset. Identifications are filtered at 1% run-specific precursor-level FDR or additionally also at 1% run-specific precursor-level eFDR (Online Methods). **(B)** Apex intensities for precursors surviving (gray) or not surviving (red) 1% run-specific precursor-level eFDR for the same data as in **(A)**, analyzed by CHIMERYS, DIA-NN and Spectronaut. **(C)** Number of fragment ions used for the quantification of precursors by CHIMERYS, DIA-NN and Spectronaut for the same data as in **(A)**. Identifications are filtered at 1% run-specific precursor-level FDR and colored by whether they also survive 1% run-specific precursor-level eFDR (Online Methods). **(D)** Example fragment ion XICs for two high-scoring precursors identified by Spectronaut (all six library fragments are shown). The top one is also identified by CHIMERYS, while the bottom one is not. The vertical dashed gray lines mark Spectronaut’s reported EG.StartRT and EG.EndRT for the corresponding precursors, with 102.36 min and 102.94 min for the top and 19.75 min and 20.05 min for the bottom XIC, respectively.

As one might expect, precursors filtered out based on eFDR have lower MS2 intensities (Figure 3B) and fewer fragment ions (Figure 3C). However, the extent to which this is observed differs substantially between the three tools, with CHIMERYS considering far more fragment ions for quantification than the other two and being more rigorous in the inclusion of fragment ions with very low intensity when using the corresponding default settings. The latter is illustrated in Figure 3D, in which the top panel shows fragment ion chromatograms for a peptide confidently identified by all three search engines, and the bottom panel shows fragment ion chromatograms for a peptide identified only by Spectronaut and for which no evidence of co-elution of fragment ions can be observed (see also Supplementary Information). Further investigations regarding the number and intensity of fragment ions, as well as the corresponding raw data (Supplementary Figure 9B-D) suggest that precursors with less than three quantifiable fragment ions with an intensity exceeding 1 or those with near-zero (or zero) intensity should be removed; either categorically or by applying stringent FDR control, which has a very similar effect (Supplementary Figure 9E). The latter brings all three software tools to a comparable level of overall identifications.

### Accurate peptide quantification from chimeric PRM and DIA spectra

One of CHIMERYS’ distinguishing concepts is its spectrum-centric processing of chimeric spectra. Apart from peptide identification, it also derives spectrum-centric quantitative information in the form of CHIMERYS coefficients, which can be interpreted as the interference-corrected total ion current for a given peptide in this MS2 spectrum (Online Methods). If none of the matched fragments for a peptide are shared with another peptide and the predicted MS2 spectrum matches perfectly to the experimental one, the coefficient is the sum of all matched fragment ions in the experimental MS2 spectrum. Hence, tracing the coefficient along retention time generates a pseudo-extracted-ion-chromatogram (XICs) that can be used to perform (relative) quantification of peptides based on their MS2 signal in PRM and DIA data. This is different from standard approaches that create XICs for (a subset of) fragment ions of a given peptide, which need to remove interfered fragment ions from quantification to maintain high precision and accuracy (Figure 4A). To assess the performance of our concept, we performed a simple PRM assay, focusing on 52 peptides from 18 human proteins spanning five orders of magnitude of cellular abundance (Online Methods). Both CHIMERYS and Skyline recovered 47 out of 52 peptides from the targeted inclusion list and CHIMERYS’ automatically-generated MS2-based quantification was in excellent agreement (R=0.99) with the manually curated values obtained from Skyline (Figure 4B). Without any additional effort, CHIMERYS identified and quantified 1,400 further peptides that were not designed to be in the assay but that happened to be co-isolated along with the targeted peptides (Supplementary Figure 10A-C). CHIMERYS effectively automates the processing of PRM data because it removes the manual curation steps often required in Skyline. These include dealing with shared fragment ions and co-isolated peptides (both used in CHIMERYS but removed in Skyline).

**Figure 4.**
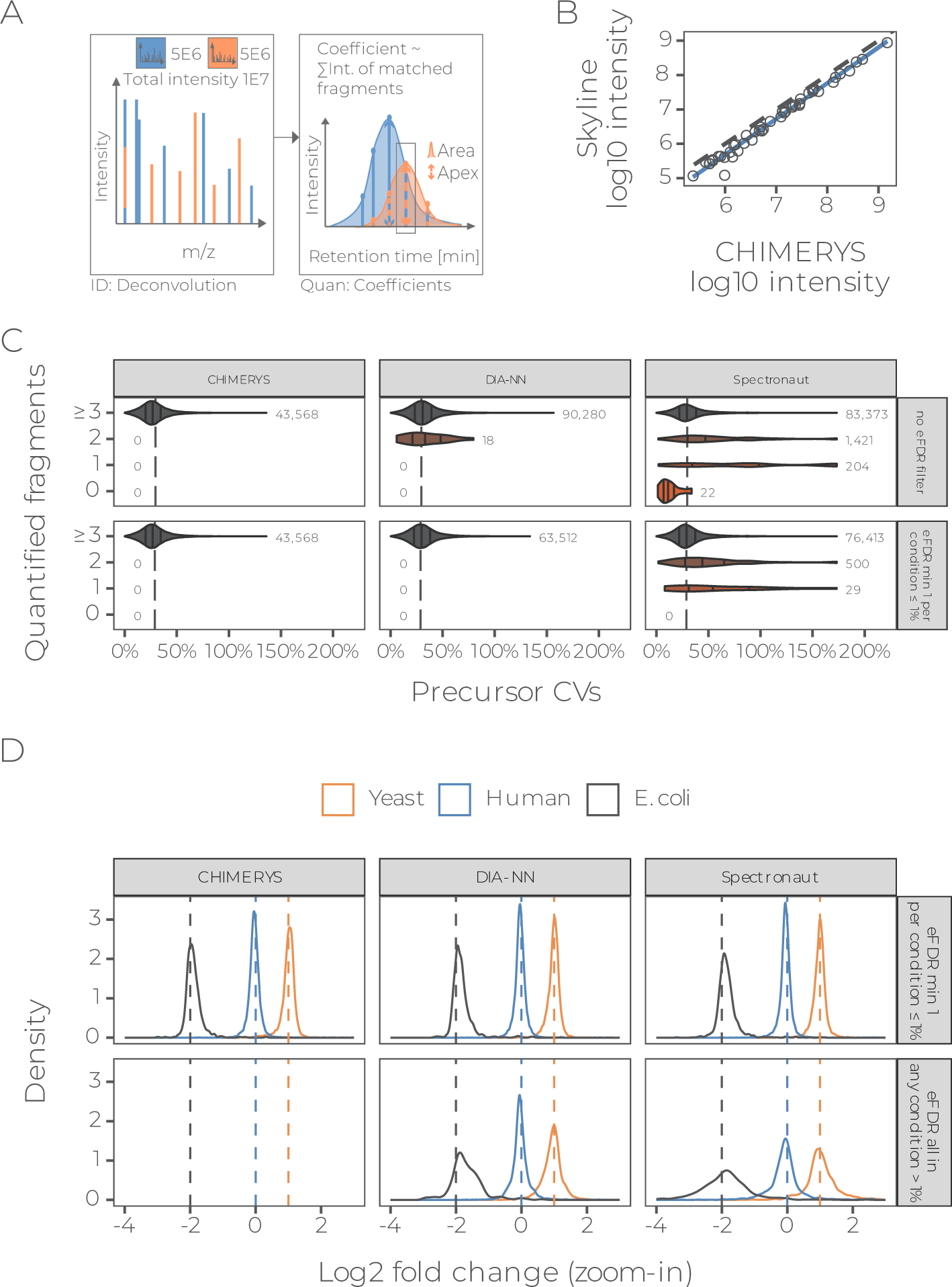
Coefficient-based quantification. **(A)** CHIMERYS quantifies precursors based on MS2 spectra by tracing their spectrum-centric coefficients – interpretable as interference-corrected TIC – over retention time, followed by apex intensity extraction or trapezoidal peak area approximation. **(B)** Correlation of CHIMERYS’ MS2-based quantification to Skyline’s MS2-based quantification on a PRM dataset targeting 52 peptides from 18 human proteins across the entire dynamic range of a human proteome. **(C)** Violin plots depicting CV distributions of precursors identified by CHIMERYS, DIA-NN or Spectronaut at 1% self-reported FDR (top) or additionally requiring at least 1 replicate per condition to also meet 1% entrapment FDR (bottom, peptide eFDR approach) at the precursor-level in the run-specific context, quantified based on 0, 1, 2, and 3 or more fragment ions. Data is from triplicate 2-hour DIA single-shot measurement from two different conditions, acquired on an Orbitrap QE HF-X with 8 Th isolation windows from the LFQbench-type dataset. Vertical dashed lines depict the median across all search engines with the respective filtering applied (29.6% for filtering at self-reported FDR and 28.9% with additional eFDR filtering). **(D)** Precursor-level log2-ratio density plots for the same data as in **(C)**, stratified by whether at least one replicate per condition survived 1% eFDR based on the peptide eFDR approach (top) or not (bottom). Replicate measurements were averaged before calculating ratios.

Next, we compared the MS2-level quantitative precision and accuracy of CHIMERYS to DIA-NN and Spectronaut on the LFQbench-type dataset^22^. To avoid differences in quantification due to different methods for determining peak integration borders, we compared the three algorithms based on their implementation of peak apex quantification. When filtering the data using eFDR as discussed above, the median quantitative precision of precursors (based on coefficient of variation, CV) was 26.9%, 29.9% and 29.1% for CHIMERYS, DIA-NN and Spectronaut, respectively (Figure 4C, bottom panel).

Similarly, analyses of precursor-level ratio distributions (Figure 4D) as a measure of quantitative accuracy for the three different search engines at eFDR were comparable (mean log_2_-ratios +/−standard deviation for *Escherichia coli*, *Homo sapiens and Saccharomyces cerevisiae* of −1.90 +/− 0.25, −0.03 +/− 0.25 and 1.00 +/− 0.29 for CHIMERYS, −1.83 +/− 0.32, - 0.04 +/− 0.23 and 0.99 +/− 0.28 for DIA-NN and −1.83 +/− 0.35, −0.04 +/− 0.31 and 1.00 +/− 0.37 for Spectronaut, respectively). The above analysis demonstrates that CHIMERYS’ spectrum-centric way of quantifying peptide precursors matches the performance of Skyline on PRM data as a gold standard in the field and extends to full-scale DIA data. It also highlights the potential of CHIMERYS for scaling PRM assays to very large numbers of peptides without the need for manual intervention.

## Head-to-head comparison of DDA and DIA data, facilitated by CHIMERYS

We showed that CHIMERYS can analyze DDA and DIA data using the same concepts for the deconvolution of chimeric spectra, which enables directly comparing the two acquisition methods on data acquired from the same sample, without the need to process the data with different software packages. As one would expect, it identified more than twice as many PSMs from DIA (8 m/z isolation windows) compared to DDA (1.3 m/z isolation windows) data acquired on an Orbitrap QE HF-X (LFQbench-type dataset; Supplementary Figure 11A). However, DDA identified 52% more peptides (and 30.3% more protein groups) compared to DIA (Figure 5A, Supplementary Figure 11B). Likely, this is due to the interplay between the AGC limit and the dynamic range in MS2 spectra, which we already observed for WWA data (see section above). In contrast, relative quantitative data completeness was higher for DIA than for DDA data when filtering for peptides that met 1% FDR in the global, but not necessarily the run-specific context and enabling ‘match between runs’ for DDA using the Minora Feature Detector in Proteome Discoverer^27^ (78% versus 55.4% of peptides quantified in two out of three replicates per condition in DIA and DDA, respectively, Figure 5A). This resulted in very similar numbers of peptides being quantified in two out of three replicates (56,322 and 52,161) for DDA and DIA, respectively.

**Figure 5.**
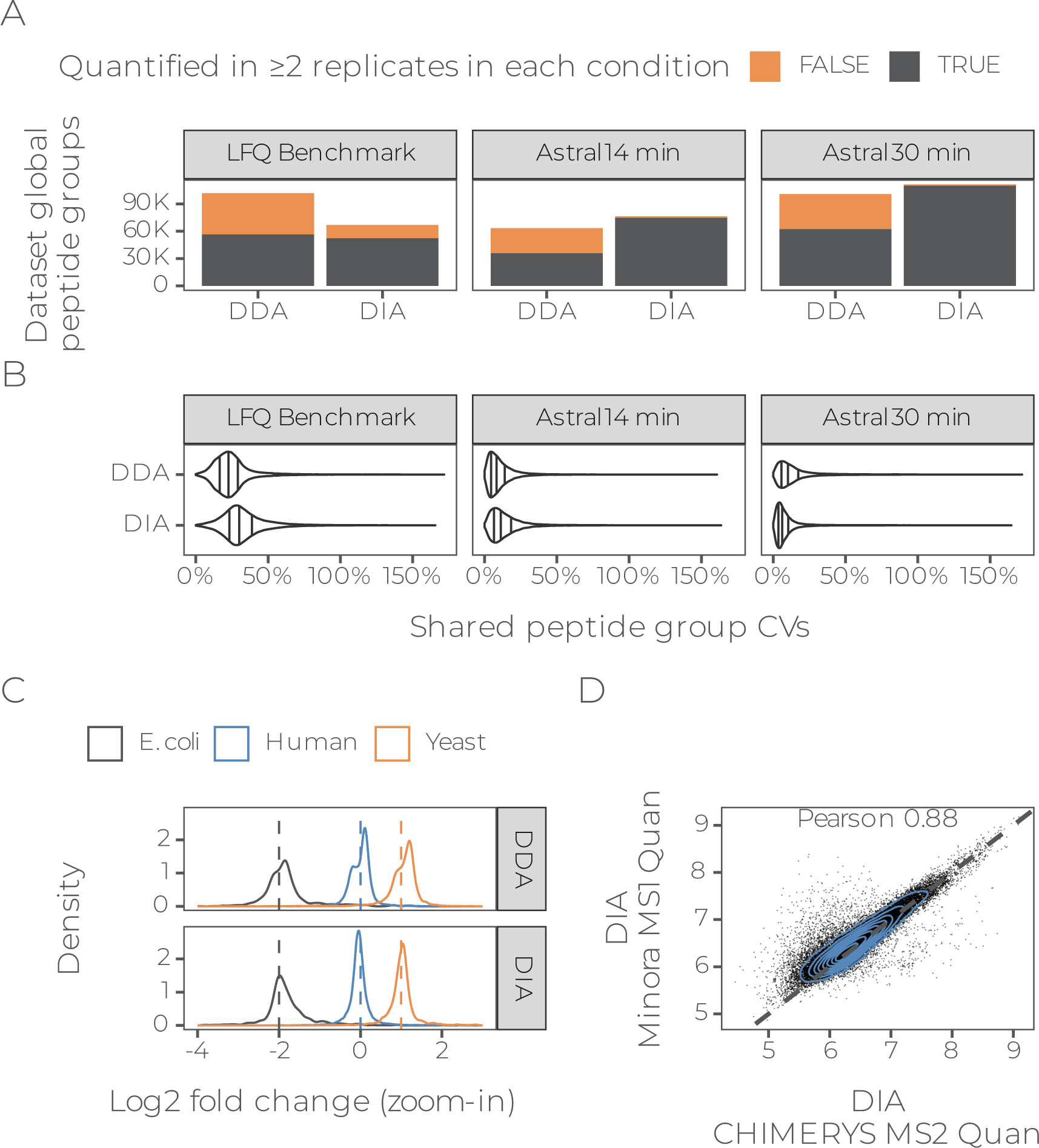
Comparing DDA and DIA data. **(A)** Peptide groups identified by CHIMERYS in triplicate 2-hour DDA (left) or DIA (right) single-shot measurement from two different conditions from the LFQbench-type dataset, acquired on an Orbitrap QE HF-X with 1.3 or 8 Th isolation windows, respectively, or in triplicate 14 min or 30 min single-shot measurements from a HeLa sample acquired in DDA (left) or DIA (right) on an Orbitrap Astral^29^. FDR was controlled at 1% at the global peptide group-level. Match between runs was used for DDA data and for DIA data, peptides were quantified irrespective of their run-specific FDR. **(B)** Peptide-level CV distributions for shared peptide groups from the same data as in **(A)**. **(C)** Peptide-level log_2_-ratio density plots for shared peptide groups from the same data from the LFQbench dataset as in **(A)**. **(D)** Scatter plot of peptide-level apex quantification between CHIMERYS’ MS2-based quantification (x-axis) and Minora’s MS1-based quantification (y-axis) for a DIA raw file from the LFQbench dataset.

Perhaps the more interesting comparison is that of DDA vs DIA using the same isolation window (here 2 m/z). This has recently become possible because modern, fast scanning instruments blur the border between DDA and DIA^28^. Interestingly, both 14 min (∼100 SPD) and 30 min (48 SPD) gradients on an Orbitrap Astral^29^ yielded similar numbers of PSM, peptide and protein group identifications for DDA and DIA (Figure 5A, Supplementary Figure 11A-B). The small differences in favor of DIA are likely due to the higher scan rate of the Orbitrap Astral in DIA mode. Again, relative quantitative data completeness was much better for DIA than for DDA (97.9% and 98.7% of peptides quantified in two out of three replicates vs 56.3% and 61.7% for the 14 min and 30 min gradients, respectively; Figure 5A). This data suggests that DIA and MS2-based quantification should be preferred over DDA and MS1-based quantification when performing label-free single-shot measurements on fast scanning instruments. Comparing CV distributions of peptides detected by DDA and DIA in the three datasets revealed that DDA was slightly more precise on the LFQbench-type dataset, while DIA was slightly more precise on the 30 min Orbitrap Astral dataset (Figure 5B). Quantitative accuracy appeared to be generally better for DIA on the LFQbench-type dataset (Figure 5C). However, closer inspection suggests that this is due to a problem with the samples rather than with MS1-based quantification *per se*, since the accuracy of MS1- and MS2-based quantification of the DIA data is comparable (Supplementary Figure 11C). In fact, CHIMERYS’ MS2-based quantification was highly correlated (R=0.88) to the MS1-based quantification implemented in Proteome Discoverer on the same raw data (Figure 5D), suggesting that the two quantification methods could be combined in the future in CHIMERYS.

## Discussion

In many ways CHIMERYS returns to the very old concept of analyzing tandem mass spectra, one at a time. At least for the task of peptide identification, this so-called spectrum-centric approach places the core analytical evidence acquired by the mass spectrometer at the center of all data analysis. This comes with a number of important advantages. First, any proteomic data type (DDA, DIA, PRM) can be treated the same and CHIMERYS is the first software implementation that stringently follows this unifying philosophy. Second, there is no principle difference between identifying a single or multiple peptides from the same MS2 spectrum and skilled scientists have done so since the early days of proteomics. The added sophistication is that artificial intelligence can predict the tandem mass spectrum of any peptide with outstanding accuracy so that it is possible to deconvolute even highly chimeric spectra by maximizing the explained intensity in an MS2 spectrum using a minimal set of peptides to do so. Third, statistical methods for PSM-level FDR control are conceptually well worked out and have reached a very high level of practical refinement, again including the use of artificial intelligence that can predict the tandem mass spectrum of any target or decoy peptide with the same accuracy, ensuring fair competition between targets and decoys. Fourth, the plausibility of an identification can be further assessed (albeit not automatically) beyond statistics by visual inspection in the context of the full MS2 spectrum and e. g. looking out for fragment ions that were not part of the deep learning model and have thus not yet been used for identification. A current limitation of CHIMERYS in this context is that peptides carrying modifications that are not yet covered by the underlying deep learning model escape detection. It can be anticipated that this limitation will diminish over time as deep learning models start to emerge that are capable of generalizing to modifications or fragmentation methods that they have not yet been trained for^30^.

Akin to other software tools, CHIMERYS also uses the information contained in the MS2 spectrum for peptide quantification. However, unlike all other DIA software, it does not set a fixed number of fragment ions to consider and instead always uses all the fragment ions that have led to an identification in a given MS2 spectrum, but in relative proportion to how much they contributed to the actual signal in the MS2 spectrum (important for the frequent case of fragment ions that are shared between peptide candidates). CHIMERYS uses the sum of these fragment ion intensities rather than the individual fragment intensities to find the apex of a chromatographic peak. This makes the overall quantification more robust against weak signals and spurious detections as encountered in e. g. single cell proteomics data. The results indicate that quantitative precision and accuracy closely match that of PRM data, which is often considered to be the gold standard for peptide quantification. In this context it is interesting to note that CHIMERYS also “en passent” automates the analysis of PRM experiments.

We consciously decided to rate data quality over quantity such that reported peptide identification and quantification results are rather conservative and other software tools may sometimes seemingly outperform CHIMERYS (see also Supplementary Information). However, when applying rigorous and consistent criteria for peptide detection and quantification, these differences diminish. A perhaps unexpected finding in this regard is that DIA data is often not nearly as complete as default processing parameters of DIA search engines report. Again, and not surprisingly, this is particularly true for low abundant samples or low abundant peptides within a sample. The reasons for this could be manifold and investigating them comprehensively goes beyond the scope of the present study. However, it is worth mentioning that the most recent generation of mass spectrometers have driven sensitivity to the point of single ion detection. As a result, MS2 spectra have at least some low level of signal at nearly every m/z. Many of these may not even stem from peptides but will create a situation in which “something” can be easily found everywhere and all the time, leading to data completeness that bears little if any actual justification. In addition, the increasing volume and density of MS-based proteomic data keeps challenging the scalability of the assumptions underlying data processing tools. Reassuringly, the community of proteomics software developers and users are increasingly aware of these recurring challenges, as it is in everybody’s best interest to ensure that software tools can be trusted and used at face value. CHIMERYS makes a valuable contribution in this context and a particularly exciting prospect is that the latest LC-MS/MS hardware along with the latest software solutions will soon overcome the historically grown divide in the field between DDA and DIA.

## Online Methods

### Deep learning for peptide property prediction

#### Data preprocessing – fragmentation prediction

Publicly available data from various PRIDE identifiers were used as a training data foundation, notably the ProtomeTool project^31^. RAW data were downloaded and – where available – MaxQuant^8^ search files were utilized. PSMs from MaxQuant’s msms.txt were merged with the unprocessed scans extracted from RAW files with Thermo Fisher’s Raw-FileReader (http://planetorbitrap.com/rawfilereader). Data were filtered using various quality criteria, e.g. Andromeda score. The top 3 ranked PSMs by Andromeda score per sequence, charge, collision energy combination were selected across all files. For the PSMs, b- and y-ions, as well as several neutral losses (e.g. water, ammonia and carbon monoxide, phosphoric acid losses) were annotated for charge states 1-3. Amino acid tokens in the peptide sequence, precursor charge, and fragmentation type were one-hot encoded, and collision energy was normalized to [0, 1]. The data contains 25M PSMs and was split into 70% training, 20% test and 10% validation sets, ensuring that peptide sequences are not shared across splits.

#### Data preprocessing – retention time prediction

The same unprocessed data was transformed similarly for retention time training with the following differences: in addition to a stringent Andromeda score threshold, only the top-ranked PSMs by Andromeda score per sequence and file, and only the 5 top-ranked PSMs per sequence across all files were retained. Retention time was z-score normalized per file to account for differing run lengths. The resulting dataset contains 5.2M PSMs split into 70% training, 20% validation, 10% test sets, ensuring that peptide sequences are not shared across splits.

#### Model architecture

INFERYS 3.0 models for fragmentation and retention time share a similar architecture. Sequences are processed with a PositionalEmbedding^32^, a Transformer block and a GRU^33^ layer (gated recurrent unit). For fragmentation prediction, additional meta information (precursor charge, collision energy and fragmentation type) are injected via a custom TransformerMixin layer to the sequence embedding outputs of the Transformer block (before the GRU layer). The TransformerMixin embeds one input parameter to the dimensionality of a given Transformer output embedding and applies the product of the two to another Transformer layer. The final Transformer embedding is projected to the task-dependent output dimension (e.g. 1 output dimension for retention time). Models are built and trained with tensorflow 2.11.1.

#### Model training

The same training procedure was used for the fragmentation and retention time model. Models are trained with the Adam optimizer^34^ on the training split with early stopping, evaluating the validation split with patience 8 and a learning rate decay with a factor of 0.2 after 4 epochs without reduced validation loss for up to 200 epochs or till convergence. The retention time model optimizes mean-absolute-error and the fragmentation model optimizes normalized spectral contrast distance^15,35^. Model hyperparameters, such as learning rate, batch size, dropout, embedding dimension, positional scaling, number of attention heads, and intermediate dimensions are optimized via Hyperband^36^. For the fragmentation model, we also optimized the order of meta parameter TransformerMixin layers via Hyperband.

#### Model capabilities

The INFERYS 3.0 fragmentation model predicts a set of fragment ions consisting of b-ions, y-ions, water-loss ions ammonia-loss ions, carbon monoxide-loss ions and phosphoric acid-loss ions in charge states 1 to 3. The exact set of ions was predetermined by selecting ions that explain the most experimental intensity in the training data set, hence some ions never or rarely observed in the training data were excluded (e.g. triply-charged y1). The fragmentation model is compatible with peptides of length 7 to 30 that can contain carbamidomethylated cysteine residues (fixed), oxidized methionine residues, phosphorylated serine, threonine or tyrosine residues, Tandem Mass Tag (TMT)-labels, TMTpro-labels and SILAC lysine4, arginine6, lysine8, and arginine10 stable isotope amino acids that were generated by either collision-induced dissociation (resonance-type CID) or higher-energy collisional dissociation (beam-type HCD). The model does not show any bias in fragmentation prediction accuracy for tryptic vs. non-tryptic analytes (data not shown). The INFERYS 3.0 retention time model can predict retention times for the same peptide classes as above. The prediction capabilities of the deep learning models effectively dictate the compatibilities of the CHIMERYS workflow.

### Brief description of the CHIMERYS algorithm

#### Setup

The CHIMERYS workflow is a cloud-native API service, orchestrated by Kubernetes on Amazon Web Services (AWS) or on-premise. The environment consists of two major components: An INFERYS prediction server^37^, which delivers predictions via gRPC requests to a CHIMERYS search algorithm instance, which matches these predictions to experimental spectra.

#### Description of the identification workflow

The CHIMERYS workflow follows the setup of classic search engines: After the *in-silico* digest of the protein database and the generation of shuffled decoy sequences using a similar logic as the mimic^14^ entrapment generator (see section below), a coarse first search is performed to identify highly confident peptides for recalibration purposes. Notably, for a group of I/L isomers, CHIMERYS only scores one representative. A fast fragment ion index implementation similar to MSFragger^38^ is used to determine a ranked list of suitable candidate peptides with isotope envelopes that (partially) overlap with the MS2 isolation window (plus tolerances). Fragment ion intensities for highly-ranking candidate peptides are predicted for each spectrum, merged against the experimental MS2 spectrum and subsequently, a set of counting-(e.g. number of matching peaks between predicted and experimental spectrum) and intensity-based scores (e.g. normalized spectral contrast angle) are calculated. Candidate peptides that fall below certain cutoff criteria are removed. For example, PSMs are required to have at least three matched fragment ions, one of which must be the base peak (most abundant peak of the prediction) and another one of which must be among the top three most intense peaks of the predicted spectrum. After the initial search, a linear discriminant analysis identifies highly confident PSMs for the calibration of optimal prediction parameters of the fragmentation model (e. g. normalized collision energy; NCE), refinement learning of the retention time model and recalibration of fragment ion m/z and match tolerances. Peptide classes with few confidently identified peptides are removed entirely from the search space (e. g. peptides of length 7 carrying two missed cleavages and two oxidized methionines). In the main search, the above-described scoring functions are executed using the optimized settings and prediction parameters, albeit now also filtering candidate peptides based on their predicted retention time. CHIMERYS uses retention time tolerance windows that would allow the identification of 99% of the peptides confidently identified in the initial search. The scoring is repeated to arrive at a set of high-scoring candidate peptides as input for the deconvolution function, where the candidate peptides simultaneously compete for experimental fragment ion intensity in one concerted step. CHIMERYS uses non-negative L1-regularized regression via the LASSO, which models the experimental spectrum as a function of the matrix of candidate peptides. The algorithm aims to explain the spectrum with the fewest number of possible candidate peptides. This is achieved by letting the different predicted spectra effectively ‘compete’ to explain the experimentally observed fragment ion intensities. The algorithm reports a coefficient for each candidate peptide representing its contribution to the experimental spectrum. A coefficient bigger than 0 indicates that this candidate peptide was used to explain the experimental spectrum. Based on the resulting coefficient, a subsequent round of intensity-based scoring is executed. Here, the coefficients of the candidate peptides can be used to predict the proportional intensity of all but one candidate peptide, add them together and subsequently subtract this sum from the actual experimental MS2 spectrum to calculate what we call a ‘shadow spectrum’, i.e. the experimental spectrum with the contributions of all interfering peptides removed. Next, the above-mentioned figures of merit are calculated based on these shadow spectra without the interference of other peaks in the spectrum. Importantly, this also works for fragment ion which are shared between candidate peptides. Candidate peptides that fail to meet certain quality criteria (e. g. minimum number of most abundant peaks shared between predicted and experimental spectrum) are filtered out. A list of all remaining target and decoy PSMs per spectrum that received a coefficient > 0 and met all quality criteria including all calculated scores is generated as input for the PSM-level error estimation in mokapot^13^.

#### FDR estimation using mokapot

For error control, the initial implementation of CHIMERYS utilized Percolator^14^ 3.0.5 to aggregate all calculated scores for all target and decoy PSMs generated in a dataset. As Percolator runtime scales poorly with large input files, we exchanged it with the Python-based reimplementation termed mokapot^13^. To ensure scalability to large input lists while controlling the compute resources, we created a public pull request to the mokapot Github repository (https://github.com/wfondrie/mokapot/pull/100) after we rewrote large parts of mokapot’s logic to allow streaming of data from disk, introduced RAM limits and implemented more performant data structures. Mokapot is executed using the following parameters: Training FDR of 1%, a training subset of 400k, and 10 iterations for training. We specifically prevent mokapot from only retaining the top-scoring PSM per spectrum. Afterwards, the resulting PSM-level q-values, support vector machine (SVM) scores and posterior error probabilities (PEPs) are attached to the corresponding PSMs. Peptides containing leucine/isoleucine (I/L) isomers in the search space are added back to the results with identical scores and are flagged as *ambiguous*.

#### MS2 quantification workflow

CHIMERYS determines raw file-specific peak apex retention times as the CHIMERYS coefficient-weighted mean of RT deltas relative to the gradient length based on PSMs meeting 1% run-specific PSM-level FDR. If an external inclusion file was used, PSMs meeting 1% run-specific PSM-level FDR including their relative retention times and CHIMERYS coefficients from the list are also considered. If no PSMs meet 1% run-specific PSM-level FDR for a given precursor in a given raw file, the apex for said precursor in this raw file is calculated using the same logic as above, but based on PSMs meeting 1% run-specific PSM-level FDR in other raw files and the inclusion file. CHIMERYS in its current implementation then estimates maximum integration borders per raw file as the 99% quantile of peak widths at base (not full width at half maximum; FWHM) from precursors with at least three PSMs surviving a run-specific PSM-level FDR threshold of 1%. These maximum integration borders are then applied to each precursor in this raw file, leading to relatively wide integration borders, particularly for low-abundant precursors. Afterwards, quantification of PRM and DIA data is performed by either trapezoidal integration of the CHIMERYS coefficients from each precursor in a set of consecutive MS2 spectra sharing the same isolation window within the integration borders, or by using the highest CHIMERYS coefficient within the integration borders as the elution peak apex intensity. One missing CHIMERYS coefficient in a series of consecutive MS2 scans with the same isolation window is allowed (gap scan) and a contribution of 0 is inserted to any further scan with missing data points, which act as boundary for peak area integration. Notably, at this point, CHIMERYS coefficients are taken from PSMs irrespective of their run-specific PSM-level FDR. However, CHIMERYS coefficients will only be used from peptide precursors that met CHIMERYS’ quality criteria (e. g. a minimum of three peaks matched between the predicted and the experimental spectrum) and are located in the vicinity of the determined peak apex. As such, at least one confidently identified PSM across all raw files is required to generate quantitative values based on PSMs around the determined peak apex in each raw file. As such, CHIMERYS will quantify precursors that fail to meet run-specific precursor-level FDR thresholds. Users are free to filter their list of precursors at 1% global precursor-level FDR (precursor was confidently identified in at least one raw file) or additionally also at 1% run-specific precursor-level FDR. The latter will reduce data completeness and is more conservative. However, we have shown that often, these quantifications are precise and accurate, so we recommend to work with precursors filtered to 1% global precursor-level FDR during exploratory data analysis and turn to run-specific precursor-level FDR for the validation of interesting hits.

#### Post-processing of CHIMERYS’ PSM-level outputs

CHIMERYS 2.7.9 is integrated into Thermo Fisher Scientific Proteome Discoverer software 3.1 (PD)^27^. Hence, PD starts CHIMERYS searches on AWS by uploading an internal format containing only MS2 spectra and some auxillary information, a fasta file and the search parameters to the CHIMERYS service, which then processes the data and generates a result file. This result file is then downloaded and post-processed by PD^27^. In this study, we used the default CHIMERYS processing and consensus workflows with minor modifications. Briefly, all DDA data processing was using the PSM-grouper node to generate peptide groups, which were then validated using qvality^39^. For DIA data, we used a special PCM-grouper node, which enables the calculation of run-specific and global precursor-level FDR. MS1-based quantification was performed using the Minora Feature Detector with default settings. MS2-based quantification was performed using the MS2 Fragment Ions Quantifier node with default settings.

### Data generation

#### External data

The following external data were downloaded from PRIDE and processed with the respective search engines. In brief, ASTRAL data extracted from Gutzman et al., 2024 (PXD046453)^29^, Arabidopsis and Halobacterium data from Müller et al., 2020 (PXD014877)^40^, body fluid data from Bian et al., 2020 (PXD015087)^41^, secretome data from Tüshaus et al., 2020 (PXD018171)^42^ and triple species mix as well as HeLa data from the LFQBench-type dataset by Van Puyvelde et al., 2022 (PXD028735)^22^. Notably, peptides carrying oxidized methionine residues were excluded from all analyses of the LFQBench-type dataset, since raw files showed differential oxidation (data not shown). An itemized mapping of external data processed as part of this study to their source and respective search settings will be made available upon publication.

#### Internal data

The following datasets were generated in house: FFPE, gradient comparison, wwDDA, instrument generations and PRM data.

#### Cell culture and sample preparation

Human HeLa and pancreatic mouse cells (ATCC, CCL-2) were cultured under standard conditions at 37°C with 5% CO_2_ in DMEM medium supplemented with 10% fetal bovine serum (FBS) and 100 U/mL penicillin (Invitrogen). At around 80% confluence cells were washed three times with PBS buffer before UREA lysis (8M Urea, 80 mM Tris pH 7.6, 1x protease inhibitor) was performed for 5 minutes on ice. Cell lysate was clarified by centrifugation (20,000 x g for 10 min).

In solution protein digest was conducted as following. First, proteins were reduced with 10 mM DTT at 37°C for 1 h, followed by alkylation with 2-chloroacetamide (CAA) at a final concentration of 55 mM for 45 min at room temperature in the dark while shaking on a thermo shaker. After the addition of five volumes of 50 mM Tris (pH 8), trypsin digest was performed overnight by adding trypsin twice (1:100) after a primary incubation time of 4h. Desalting was performed using Sep-Pak columns according to the user manual. Human brain FFPE samples were digested using a SDS lysis protocol followed by digestion with the SP3 approach as described in detail in Tüshaus et al. 2023^42^.

#### LC-MS/MS

FFPE, gradient comparison and wwDDA data were acquired on a micro-flow LC coupled via a HESI source to an Q Exactive HF-X hybrid quadrupole-Orbitrap mass spectrometer (Thermo Scientific). Optimization of the micro-flow LC setup as well as technical details were previously published in Bian et al., 2020^41^. In brief, peptide separation was performed on an Acclaim™ PepMap™ 100 C18-HPLC-column (15 cm lengths, 1 mm inner diameter, 2 µm particle size, #164711, Thermo Fisher Scientific) at 55°C. Linear gradients with buffer A (0.1% FA, 3% DMSO in dH_2_0) and buffer B (0.1% FA, 3% DMSO in ACN) from 3 to 28% B were run at 50 µL/min. Sample loading, column wash and equilibration was performed at 100 µL/min. Source settings were applied as following: 320°C capillary temperature, 3.5 kV spray voltage, 300°C auxiliary gas. MS data were acquired at a normalized collision energy of 28 %, in Top20 mode, at an m/z range of 360–1300, AGC target of 3E6 and 1E5, maximal injection time of 50 ms and 22 ms, resolution of 60 k and 15 k on MS1 and MS2 level, respectively. The MS2 isolation window width was 1.4 Th in standard DDA runs and increased up to 20.4 Th for wide window acquisition DDA as indicated in the figure legends.

Ion trap data were acquired with an Orbitrap Eclipse™ Tribrid™ mass spectrometer (Thermo Scientific) that was coupled to a Dionex UltiMate 3000 RSLCnano System (Thermo Scientific). Samples were transferred onto a trap column (75 μm x 2 cm, 5 μm C18 resin Reprosil PUR AQ - Dr. Maisch). After washing with the trap washing solvent (5 μl/min, 10 min), samples were separated on an analytical column (75 μm x 48 cm, 3 μm C18 resin Reprosil PUR AQ - Dr. Maisch). A 70 min method, including a 50 min gradient, was performed with a flow rate of 300 nl/min (4 % B up to 32 % B within 50 min). Solvent A: 0.1 %v FA, 5%v DMSO in dH_2_O, Solvent B 0.1 %v FA, 5 %v DMSO in ACN. MS1 scans were acquired with an Orbitrap resolution of 60 k, within a scan range of 360-1300 m/z, a maximum injection time of 50 ms, a normalized AGC target of 100% and RF lens of 40 %, including charge stage 2 to 6, with an exclusion time of 25 s. MS2 scans were performed with the ion trap with a normalized AGC target of 200%, a maximum injection time of 25 ms and either with an HCD collision energy of 31% (wwDDA) or with an CID collision energy of 35 % (CID). Quadrupole isolation window was varied between 0.4, up to 5.0 m/z as indicated in the figure.

Data for the instrument comparison were assembled from 1-hour HeLa QC runs, acquired over several years at the Chair of Proteomics and Bioanalytics at the TUM. They were run on various liquid chromatography systems, employed diverse instrument-specific settings, slightly different gradients and used different batches of HeLa digest, prepared in-house.

#### Targeted assay generation

A simple PRM assay was devised by randomly selecting 18 proteins and 2-3 peptide precursors each across the whole measured intensity range from a 1-hour HeLa run analyzed on a Orbitrap Fusion Lumos mass spectrometer (Thermo Scientific). A total of 51 precursors were put into an inclusion list in addition to 14 precursors corresponding to a retention time standard. A 1-hour HeLa sample was analyzed in PRM mode: MS2 spectra were acquired using 0.4 Th isolation window, a maximum injection time of 100 ms, HCD collision at a normalized collision energy of 28 % and read out in the Orbitrap at 15k resolution.

### Data processing and evaluation

#### Data analysis using various search algorithms

A variety of search engines have been applied in the scope of this study. Unless otherwise noted, default settings were used including trypsin digestion (with proline rule, so cutting after K/R, but not before P), carbamidomethylation of cysteine as fixed modification as well as methionine oxidation as variable modification. Peptide lengths were limited to 7-30 for comparisons between CHIMERYS, DIA-NN and Spectronaut. All searches were performed against the canonical FASTA files of the corresponding species or species mix. A contaminants FASTA was utilized in all searches to control for contaminations^43^. Entrapment searches were performed as described in detail its own section. All FASTA files used in this study will be made publicly available via PRIDE upon publication.

##### CHIMERYS

Searches were performed using CHIMERYS v2.7.9 from PD v3.1.0.622 using default settings. For quantification, non-normalized intensities from Minora Feature Detector were used. PSM, precursor, peptide group and protein group IDs were extracted from .pdResult files after the removal of decoys and PSM-, precursor-, peptide group- or protein-level q-value filtering. For ion trap data, an additional TopN Peaks filter node was used before the CHIMERYS node with TopN set to 15 and mass window to 100 Da.

##### Sequest HT

Searches were performed from PD v3.1.0.622 using default settings. For quantification, non-normalized intensities from Minora Feature Detector were used. PSM and peptide group IDs were extracted from .pdResult files after the removal of decoys and PSM- or peptide group-level q-value filtering.

##### MS Amanda

Seaches were performed using MS Amanda v2.0 from PD v3.0.0.201 using default settings. For quantification, non-normalized intensities from Minora Feature Detector were used. PSM and peptide group IDs were extracted from .pdResult files after the removal of decoys and PSM- or peptide group-level q-value filtering.

##### Comet

Searches were performed from PD v3.1.0.622 using default settings. For quantification, non-normalized intensities from Minora Feature Detector were used. PSM and peptide group IDs were extracted from .pdResult files after the removal of decoys and PSM- or peptide group-level q-value filtering.

##### MSFragger

Searches were performed using MSFragger v20.0 using the “Default” workflow. Additionally, MSFragger v21.1 was utilized with the “WWA” workflow and “DDA+” data type, which considers candidate peptides in the full MS1 isolation window and reports up to the top five PSMs instead of only one. Precursor-level IDs were extracted from ion.tsv files, which were already filtered for decoys and precursor FDR. Peptide group-level IDs were rolled up from precursor-level data, controlled at 1% precursor-level FDR.

##### MetaMorpheus

Searches were performed using MetaMorpheus v0.0.320 with default settings. PSM-level IDs were extracted from AllPSMs.psmtsv files after the removal of decoys and PSM-level q-value filtering. MetaMorpheus does not feature peptide group-level IDs, so these were rolled up from the PSM level, controlled at 1% PSM-level FDR.

##### MS-GF+

Searches were performed using MS-GF+ v2022.04.18 with default settings. The output .mzid file was converted to .tsv, from which PSM-level IDs were extracted after the removal of decoys and PSM-level q-value filtering. MS-GF+ does not feature peptide group-level IDs, so these were rolled up from the PSM level, controlled at 1% PSM-level FDR.

##### MaxQuant

Searches were performed with default settings in MaxQuant v2.4.2.0. Peptide group IDs were extracted from modificationSpecificPeptides.txt files after filtering for decoys, which were already peptide group-level FDR filtered.

##### DIA-NN

Library-free searches were performed using DIA-NN 1.8.1^23^. Mixed species samples were analyzed with a classic, concatenated, and peptide eFDR approach. Entrapment fastas were based on a database for the three species and a database containing contaminants. Detailed information on entrapment database generation can be found under ‘Entrapment database construction via mimic’. For all searches, spectral libraries were generated from fasta files using DIA-NN’s prediction capabilities. Default settings were used with the following adjustments: Missed cleavages = 0, Maximum number of variable modifications = 2, Quantification strategy = Peak height. ‘Ox(M)’ was added as a variable modification. Note that ‘MBR’ is checked per default for library-free searches. The output results from the report.tsv were used. For run-specific precursor filtering, results were filtered for Q.Value ≤ 0.01. Precursor.Quantity was used as precursor quantification values. Fragments for identification were counted considering fragments with Fragment.Quant.Corrected > 0. Fragments for quantification were counted by identifying fragments, for which the sum of Fragment.Quant.Corrected corresponded to Precursor.Quantity. Samples with different species composition were analyzed together.

##### Spectronaut

Raw files were analyzed using Spectronaut v18 (Biognosys) with a library-free approach (directDIA+). Mixed species samples were analyzed with a classic, concatenated, and peptide eFDR approach. Entrapment fastas were based on a database for the three species and a database containing contaminants. Detailed information on entrapment database generation can be found under ‘Entrapment database construction via mimic’. Default settings were used with the following adjustments: Max Peptide Length = 30, Missed Cleavages = 0, Max Variable Modifications = 2, Quantity Type = Height. Carbamidomethyl (C) was set as a fixed modification, and Oxidation (M) was set as a variable modification. Cross-run normalization was disabled. Samples with different species composition were analyzed together. For run-specific precursor filtering, results were filtered for EG.Qvalue ≤ 0.01. ‘EG.TotalQuantity (Settings)’ was used as precursor quantification values. Fragments for identification were counted considering fragments with F.PeakArea > 1. Fragments for quantification were counted by identifying fragments with F.PeakArea > 1 and F.ExcludedFromQuantification = FALSE.

### Post-hoc filtering of Spectronaut output files

We show that Spectronaut claims to confidently identify precursors with no apparent signal in the corresponding MS2 spectra. In order to remove these precursors, an unfiltered, fragment-level export needs to be created from within Spectronaut. Based on this report, fragments with F.PeakArea > 1 and F.ExcludedFromQuantification = FALSE can be identified. In Spectronaut, the desired number of fragments for quantification can be set (under DIA Analysis > Quantification > Interference Correction > MS2 Min). Per default, ‘MS2 Min’ is three. Hence, in this study, precursors were filtered to only those with at least ‘MS2 Min’ fragments that have F.PeakArea > 1 and F.ExcludedFromQuantification = FALSE.

### Calculation of shared fragment ions

Peptide fragment intensity predictions of PSMs identified by CHIMERYS v2.7.9 were generated using INFERYS 3.0.0. The R package rawrr^44^ was used to extract centroided m/z and intensity values from raw files. Different PSMs within the same MS2 scans were tested for shared fragment ions using two methods: directly matching predicted fragment ions (raw file independent) and checking if predicted fragment ions matched to the same raw file peaks with a 20 ppm tolerance. The fraction of shared fragment ions among all fragment ions was determined and analyzed based on amino acid position and 200 m/z bins.

### Entrapment database construction via mimic

We generated same-organism entrapment proteins in nine-fold excess relative to the number of target proteins by shuffling sequences at the peptide level using mimic^14^ with the command line flags --prepend, --empiric, --replaceI and --mult-factor 9. The goal is to generate nine different entrapment peptides per target peptide, which are isobaric, shuffled versions of it. Entrapment peptides shall have the same peptide N- and C-termini as their target counterpart and shall be as close in amino acid composition to it as possible. Briefly, mimic achieves this by first digesting the target protein database into fully tryptic peptides without missed cleavages (no proline rule, i. e. cleaving proteins after each K/R). Afterwards, the sequence of each unique target peptide is permuted randomly while keeping the termini fixed. In case permutation of a peptide generates a target or previously-generated entrapment peptide, shuffling is repeated 1,000 times. For this comparison, isoleucine and leucine are considered to be identical amino acids. If no entrapment peptide could be generated after 1,000 rounds of shuffling, amino acids are mutated. Briefly, a random amino acid is mutated to a different one while respecting the amino acid frequencies of the target protein database. Isoleucine and leucine are never mutated when using the --replaceI flag. Notably, it is ensured that target peptides with the same peptide sequence generate the same entrapment peptides. Subsequently, entrapment peptides are assembled to entrapment proteins, which are then either appended to the target database in the case of the classic fasta concatenation approach (classic eFDR) or concatenated to the corresponding target protein sequences – separated by lysine residues – in the sequence concatenation approach (concatenated eFDR). The final digested fasta approach (peptide eFDR) used a peptide-level fasta, which was generated by digesting the classic eFDR fasta file with Protein Digestion Simulator v2.4.7993.32903 before passing it to the search engine (fully tryptic digest with no missed cleavages and no proline rule). Notably, protein-C-terminal peptides cannot be identified with the concatenated eFDR approach and neither can protein N-terminal peptides with methionine excision from entrapment proteins. Mimic is available at https://github.com/percolator/mimic. A simplified web-version of it is available at https://mimicerys.msaid.io/.

### FDR definitions

We adhere to previously-established definitions of FDR in the run-specific and global context for any identification level (e. g. precursors, modified peptides, peptides, peptide groups, protein groups)^25^. For example, peptide-level FDR in the run-specific context can answer the question “Which peptides were detected at 1% FDR in this specific LC-MS/MS run?”. Peptide-level FDR in the global context can instead answer the question “Which peptides were detected at 1% FDR in at least one LC-MS/MS run of a given experiment containing multiple LC-MS/MS runs?”.

### Calculation of entrapment FDR

We empirically validated the self-reported FDR by CHIMERYS, DIA-NN and Spectronaut using entrapment experiments, which are sometimes also called double-decoy experiments. The general idea is that most search engines are black boxes that report lists of identifications at self-reported FDR. Entrapment experiments try to empirically validate this self-reported FDR.

Briefly, most if not all search engines try to control the FDR of their identifications at some percentage (usually 1 %). In practical terms, this means they try to limit for example the number of random false target PSMs in the final list of PSMs, i. e. the FDR at the PSM level. However, because the identity of random false target identifications is not know *a priori*, search engines model the behavior of random false targets with the help of decoys. Decoys are peptides that resemble target peptides, but are assumed to be absent from the sample under investigation. Many if not all search engines generate their own decoy peptides using one of several approaches (e. g. reversing protein sequences^8^). Experimental MS2 spectra are then compared to theoretical, predicted or library spectra from target and decoy peptides, which results in a score for each PSM that measures how closely they match. In classic target decoy competition, only the top-scoring PSM of each spectrum is retained. The remaining PSMs are then sorted in descending order by their score (assuming that a high score is associated with a good match) and q-values are calculated as the cumulative sum of decoys divided by the cumulative sum of targets for each PSM. Usually, these q-values are then monotonized by sorting the PSMs by their score in ascending order and calculating the cumulative minimum of the q-values for each PSM to arrive at the final q-values. Removing PSMs with a q-value > 0.01 would filter the list of PSMs to 1% FDR. Notably, as mentioned above, this approach assumes that decoys resemble random false targets and consequently that the score distribution of decoy peptides is identical in shape and magnitude to the score distribution of random false target peptides. If this is not the case, FDR could be over- or underestimated.

Entrapment or double-decoy experiments measure empirical FDR by following very similar principles as the target/decoy approach described above. Before passing the target protein database to the search engine, which then generates its own decoys (one for every peptide after *in silico* digestion of the target protein database), so-called entrapment proteins are added to it. Entrapment proteins consist of entrapment peptides, which are very similar to decoy peptides in that they should resemble target peptides, but are assumed to be absent from the sample under investigation. However, the search engine does not know which peptide is a target peptide and which peptide is an entrapment peptide. As such, it generates decoy peptides for each target and each entrapment peptide. We will call them ‘normal decoys’ and ‘entrapment decoys’, respectively. Experimental MS2 spectra are then compared to theoretical, predicted or library spectra from target, normal decoy, entrapment and entrapment decoy peptides. Now the search engine calculates its FDR as described above after sorting PSMs based on their score in descending order by dividing the cumulative sum of decoy identifications (normal decoys + entrapment decoys) by the cumulative sum of target identifications (targets + entrapments), followed by q-value monotonization. In addition, since the identity of entrapments is known to the researcher, an entrapment FDR can be calculated. This is done by sorting PSMs in descending order by their score and calculating entrapment q-values as the cumulative sum of normal decoys plus entrapments, divided by the cumulative sum of targets plus entrapments for each PSM, followed by entrapment q-value monotonization. For entrapment analyses with DIA-NN and Spectronaut, the formula for calculating entrapment FDR was slightly modified, because their final results violate the assumption that the score distribution of decoy peptides is identical in shape and magnitude to the score distribution of random false target peptides. This is because they perform two searches of the data; one to generate a filtered spectral library and a second one that uses this spectral library to analyze the same data again in a peptide-centric fashion. The filtering of the spectral library is done based on an initial FDR calculation, which will remove a substantial fraction of random false target peptides, causing the distribution of decoy peptides in the second search to not be identical anymore in shape and magnitude to the score distribution of random false target peptides. Hence, the cumulative sum of normal decoys was removed from the entrapment FDR calculation for DIA-NN and Spectronaut. As such, entrapment FDR was calculated by sorting precursors in descending order by their score and calculating entrapment q-values as the cumulative sum of entrapments, divided by the cumulative sum of targets plus entrapments for each precursor, followed by entrapment q-value monotonization. Removing PSMs with an entrapment q-value > 0.01 should also filter the list of PSMs or precursors to 1% FDR if the search engine does not have a bias in its scoring function. In other words, if there is no bias, then q-values and entrapment q-values should follow the diagonale if they are plotted against one another in a scatter plot.

Given the entrapment FDR calculations above, it is clear why we use entrapments in at least nine-fold excess relative to targets. The reason is because the search engine generates decoys for each target and each entrapment. In other words, using the same number of entrapments as targets would result in a search space consisting of one part targets, two parts decoys (one part normal decoys and one part entrapment decoys) and one part entrapments. Given the nature of random false identifications, they are equally likely to map to false targets, decoys or entrapments. However, we cannot identify them as false matches if they map to false targets. Consequently, the likelihood of a random false identification that we can identify to map to decoys is twice as high (66%) as the likelihood of it mapping to an entrapment (33%) if the number of targets and entrapments was the same. As we increase the number of entrapments that are added to the target protein database, the likelihood of a random false identification to map to decoys or entrapments becomes more and more similar. At a nine-fold excess of entrapments relative to targets, the likelihood of a random false identification that we can identify to map to a decoy is merely ∼6% higher (53%) than the likelihood of it mapping to an entrapment (47%).

Entrapments can be added to the target protein database in various different ways (see section ‘Entrapment database construction via mimic’). In the case of the concatenated eFDR approach, we had to modify the calculation of eFDR slightly for CHIMERYS, because it is no longer possible to distinguish between normal decoys and entrapment decoys. This is because they are generated internally by CHIMERYS and are both annotated with the target protein identifier, which can be used in the other entrapment approaches to identify normal decoys and entrapment decoys. Hence, for the concatenated eFDR approach, eFDR calculation is done by sorting PSMs in descending order by their score and calculating entrapment q-values as the cumulative sum of decoys plus entrapments, divided by the cumulative sum of targets plus entrapments for each PSM, followed by entrapment q-value monotonization. No change was made to the calculation of eFDR based on the concatenated eFDR approach for DIA-NN and Spectronaut.

Throughout the manuscript, we performed several entrapment analyses for CHIMERYS, DIA-NN and Spectronaut. In Figure 1B and Supplementary Figure 8, we directly compared the self-reported run-specific FDR at a given identification level (e. g. peptides or precursors) to the corresponding entrapment FDR. In Figures 3 and 4, as well as in Supplementary Figure 9, we filtered identifications either at the self-reported run-specific FDR or additionally also at the corresponding entrapment FDR. We then compared the characteristics of the identifications that only survive the self-reported FDR or also survive the entrapment FDR (e. g. precision and accuracy of quantification). For CHIMERYS, we used the q-values and SVM scores reported by qvality in PD, which either validated peptide groups in the global context (Figure 1B) or precursors in the run-specific context (all other visualizations). For Spectronaut, we used the EG.Qvalue and the EG.Cscore columns of unfiltered Spectronaut exports, which validated precursors in the run-specific context. For DIA-NN, we used the Q.Value and CScore columns of the main report.tsv, which validated precursors in the run-specific context. Unfortunately, DIA-NN and Spectronaut do not report q-values for decoy identifications and the former always filters its report.tsv to 1% run-specific precursor-level FDR. They do however report scores for decoy identifications. In order to convert these scores into q-values for the plots in Supplementary Figure 8, we interpolated the q-values given the relationship of score and q-value for target precursors in each raw file separately.

### PRM Analysis in CHIMERYS and Skyline

PRM data were searched using CHIMERYS against a Human Swissport database with default settings. The resulting MS2-based quantities and coefficient traces were extracted from the .pdresult files using custom R scripts. The .raw file was also analyzed in Skyline^9^ v22.2 using a Prosit-predicted spectral library^15^ as integrated in Skyline. Peak boundaries were manually refined for all precursors and the quantitative values for the area beneath the five most abundant fragment ions was aggregated into a quantitative measure. Correlations were calculated and visualized using custom R scripts.

### Comparison of DDA and DIA data using CHIMERYS

Unless otherwise noted, DDA and DIA files were searched with CHIMERYS v2.7.9 from PD v3.1.0.622, using Minora for MS1-level and CHIMERYS for MS2-level quantification, respectively. PSMs were filtered at 1% run-specific FDR, peptide groups and protein groups at 1% dataset global peptide group and protein FDR, respectively. Peptide groups and protein groups with FoundinSamples = 0 were discarded. Protein groups were further filtered for PsmCount > 0 and IsMasterProtein = 0. For quantification of conditions / ratios, at least 2 non-normalized intensity values > 0 were required per condition / in both conditions, respectively (or 3 if stated “all replicates”). For calculating CVs, at least 3 non-normalized intensity values > 0 were required per condition. 2D density estimates were calculated using kde2d from the R package MASS. Entrapment FDR was calculated as described above using the peptide eFDR approach.

## Abbreviations

ACN: Acetonitrile
AGC: Automatic gain control
AWS: Amazon Web Services
CAA: Chloroacetamide
CID: Collision induced dissociation; synonym for resonance-type CID
CV: Coefficient of variation
DDA: Data-dependent acquisition
DMSO: Dimethyl sulfoxide
DIA: Data-independent acquisition
FDR: False discovery rate
eFDR: Entrapment false discovery rate
FA: Formic acid
FBS: Fetal bovine serum
FFPE: Formalin-fixed paraffin-embedded tissue
FTMS: Fourier-transform mass spectrometry
FWHM: Full width at half maximum
GRU: Gated recurrent unit
HCD: Higher-energy collisional dissociation; synonym for beam-type CID
ITMS: Ion-trap mass spectrometry
I/L isomers: Leucine/Isoleucine isomers
MS1: Precursor mass spectrum
MS2: Tandem mass spectrum
m/z: Mass over charge
PBS: Phosphat buffer saline
PRM: Parallel reaction monitoring
PSM: Peptide spectrum match
PD: Proteome Discoverer Software
RT: Retention time
SILAC: Stable isotope labeling by amino acids in cell culture
SPD: Samples per day
SVM: Support vector machine
Th: Thomson, unit
TMT, TMTpro: Tandem mass tags
WWA: Wide window Acquisition, synonym for wide window DDA (wwDDA)
wwDDA: Wide window DDA, synonym for wide window acquisition (WWA)
XIC: Extracted ion chromatogram

## Data Availability

### External raw data

An itemized mapping of external data processed as part of this study to their source and respective search settings will be made available upon publication.

### Generated data

The generated mass spectrometric raw and search data of internal datasets from this study will be made available via PRIDE^45^ upon publication.

### Open-source software

The mokapot version used in this study is available as a public pull request on GitHub (https://github.com/wfondrie/mokapot/pull/100). The modifications to the mimic entrapment database generator are available on GitHub (https://github.com/percolator/mimic/). A web-version of the mimic tool can be found at https://mimicerys.msaid.io/.

### Data analysis and plotting scripts

The custom R scripts underlying the data analysis are available upon request.

## Acknowledgements

The authors wish to thank numerous scientific colleagues for their input, discussions, and support. The authors want to expressively thank the Proteome Discoverer (PD) software development team at Thermo Fisher Scientific for their collaboration, support, and contributions on the successful integration of CHIMERYS into PD and the scientific discourse on the results. The authors wish to thank Elmar Zander for consulting on mathematical topics. The authors wish to thank Matthew The for consulting on entrapment experiments and FDR control. The authors also wish to thank their colleagues Dulguun Bold, Jeremiah Santoso, Agnes Guevende and Mohammed Al Kiddeh for various contributions to the software.

## Author contributions

M.F. and M.W. conceived the study. M.F., M.W., and D.P.Z. developed and evaluated the initial prototype. F.S., P.S., T.S., M.G., I.B., S.B.F. and S.G. developed, implemented, and optimized the algorithms. S.G., V.S., S.B.F., L.M. and M.G. developed the deep learning models. T.S., M.G. and F.S. orchestrated the implementation of software modules and the deployment of the software. M.F., D.P.Z., F.S., M.G., M.T.B. and A.H. evaluated the algorithm. M.F., M.T.B., J.T., A.H., and D.P.Z. processed the result data. M.F., M.T.B., J.T, A.H. and D.P.Z. performed the data analysis. L.E. helped in the preparation of the Figures. M.F., D.P.Z., M.B., A.H., F.S., P.S., S.G., T.S., J.T., B.K. and M.W. provided critical feedback, discussed the results, and consulted in revisions. M.F., D.P.Z., J.T., B.K. and M.W. wrote the manuscript.

---

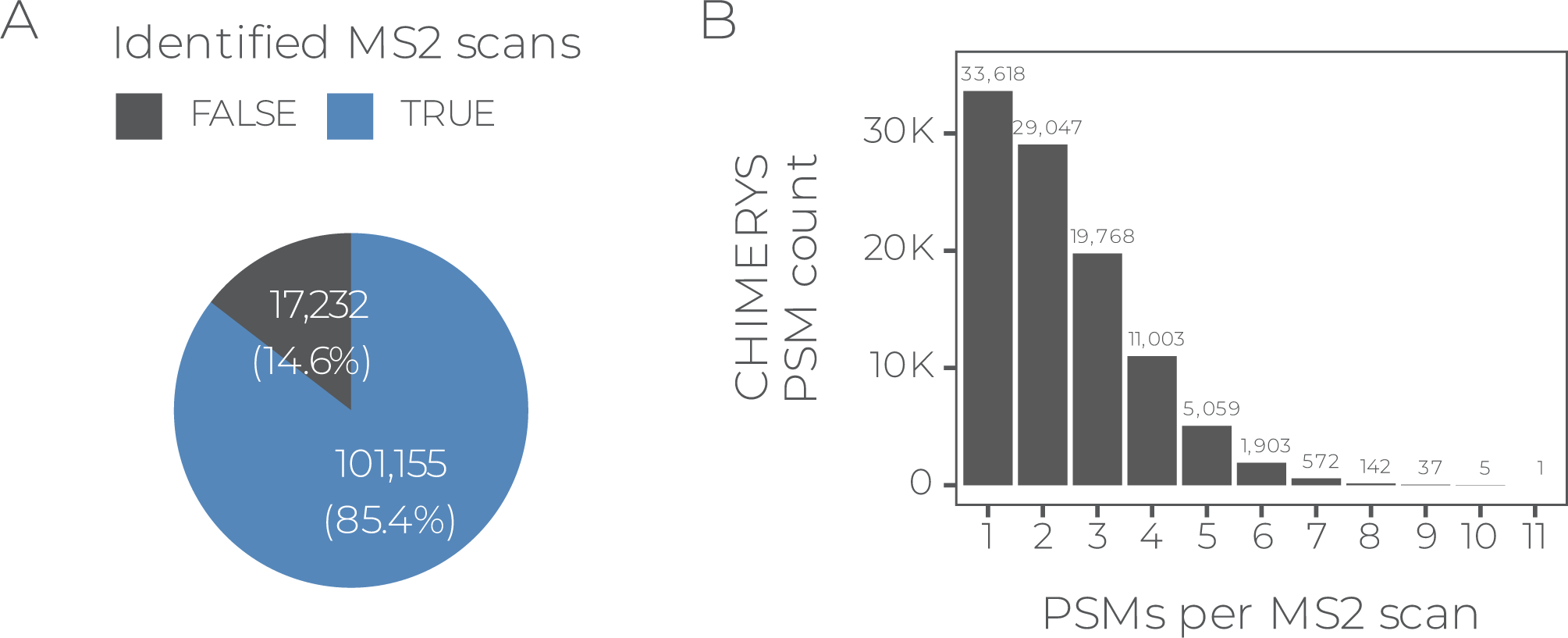

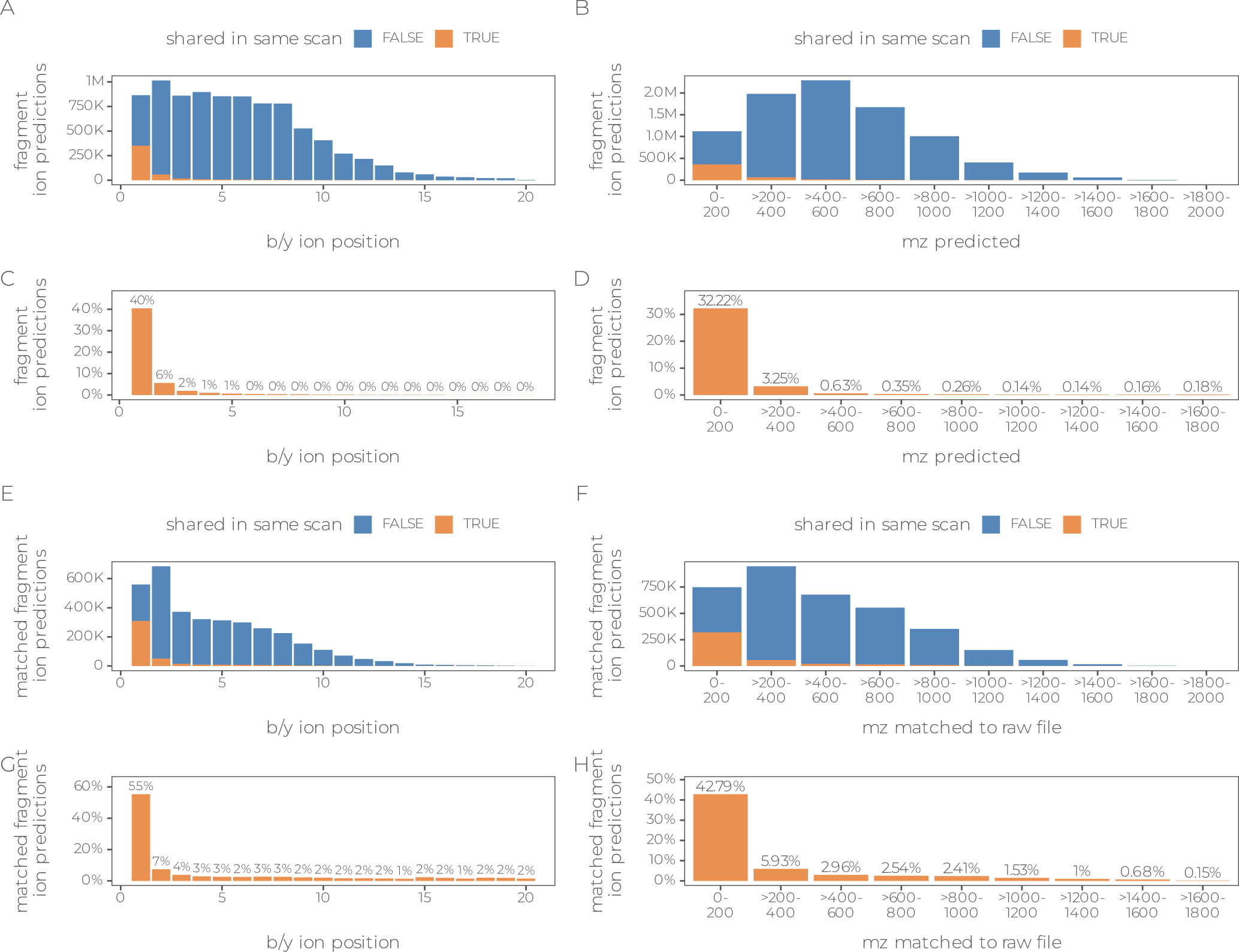

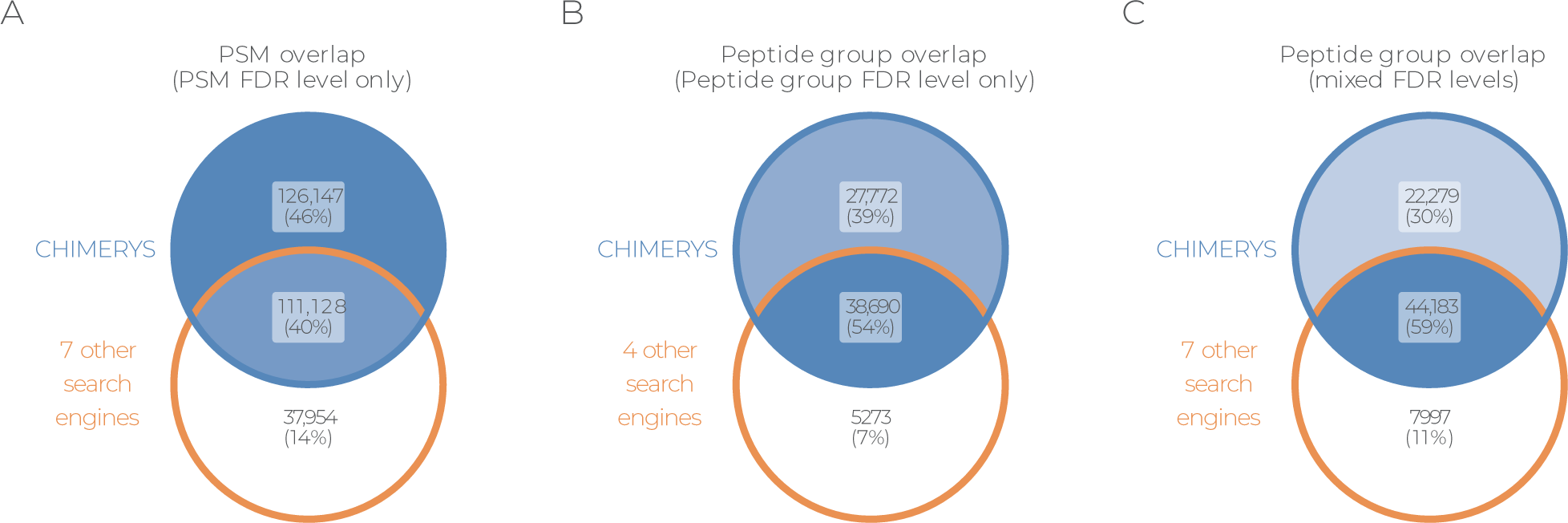

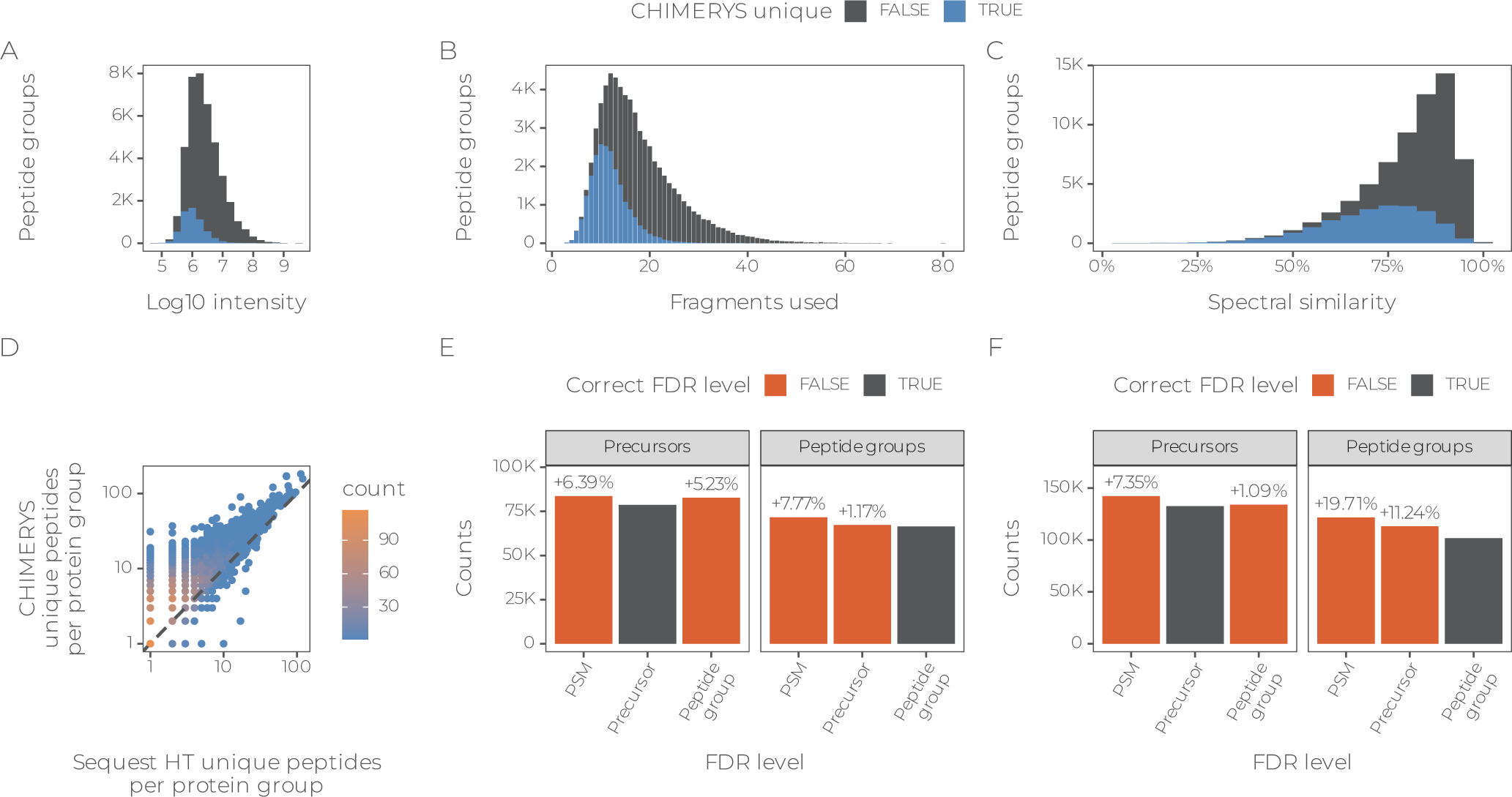

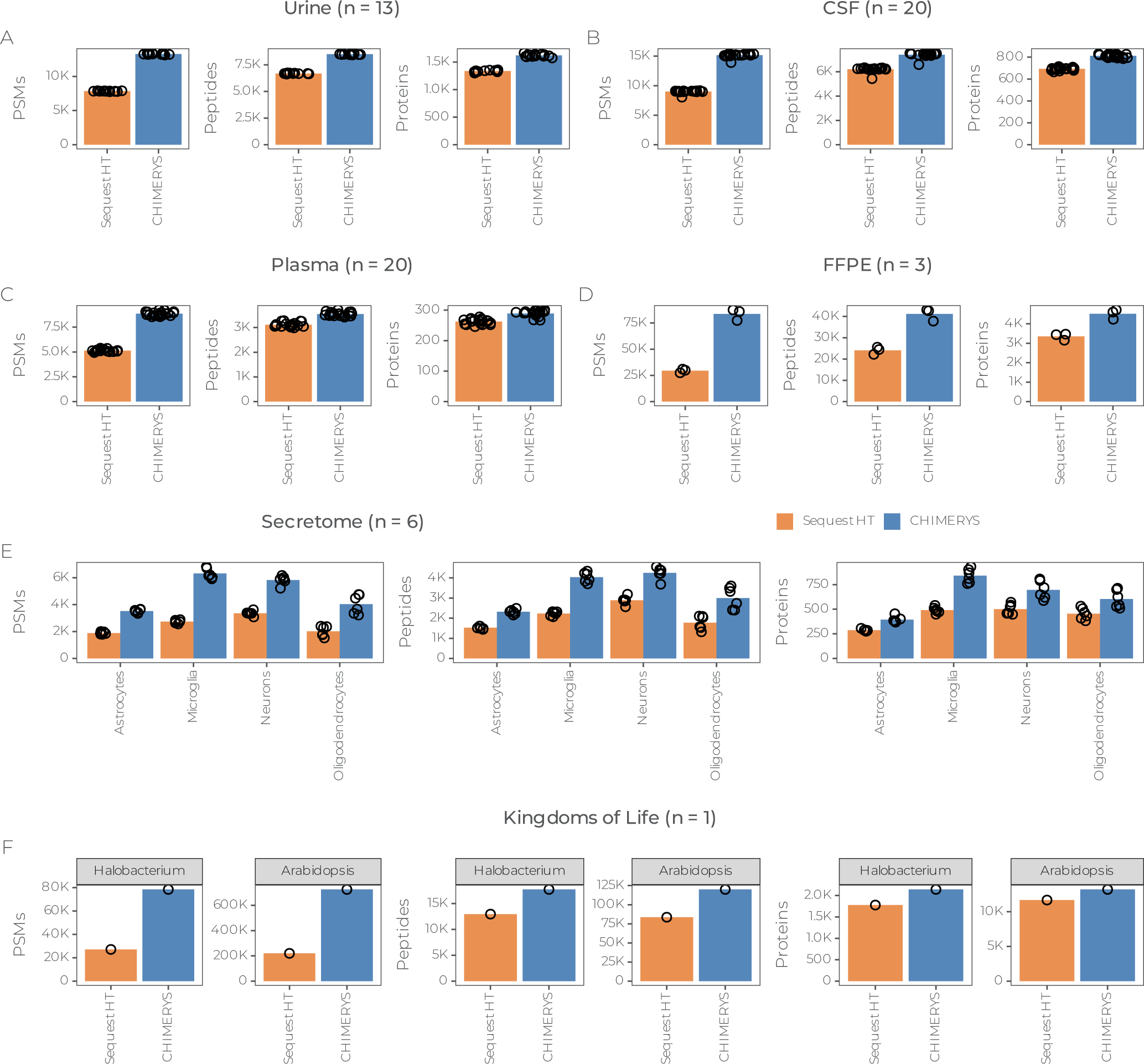

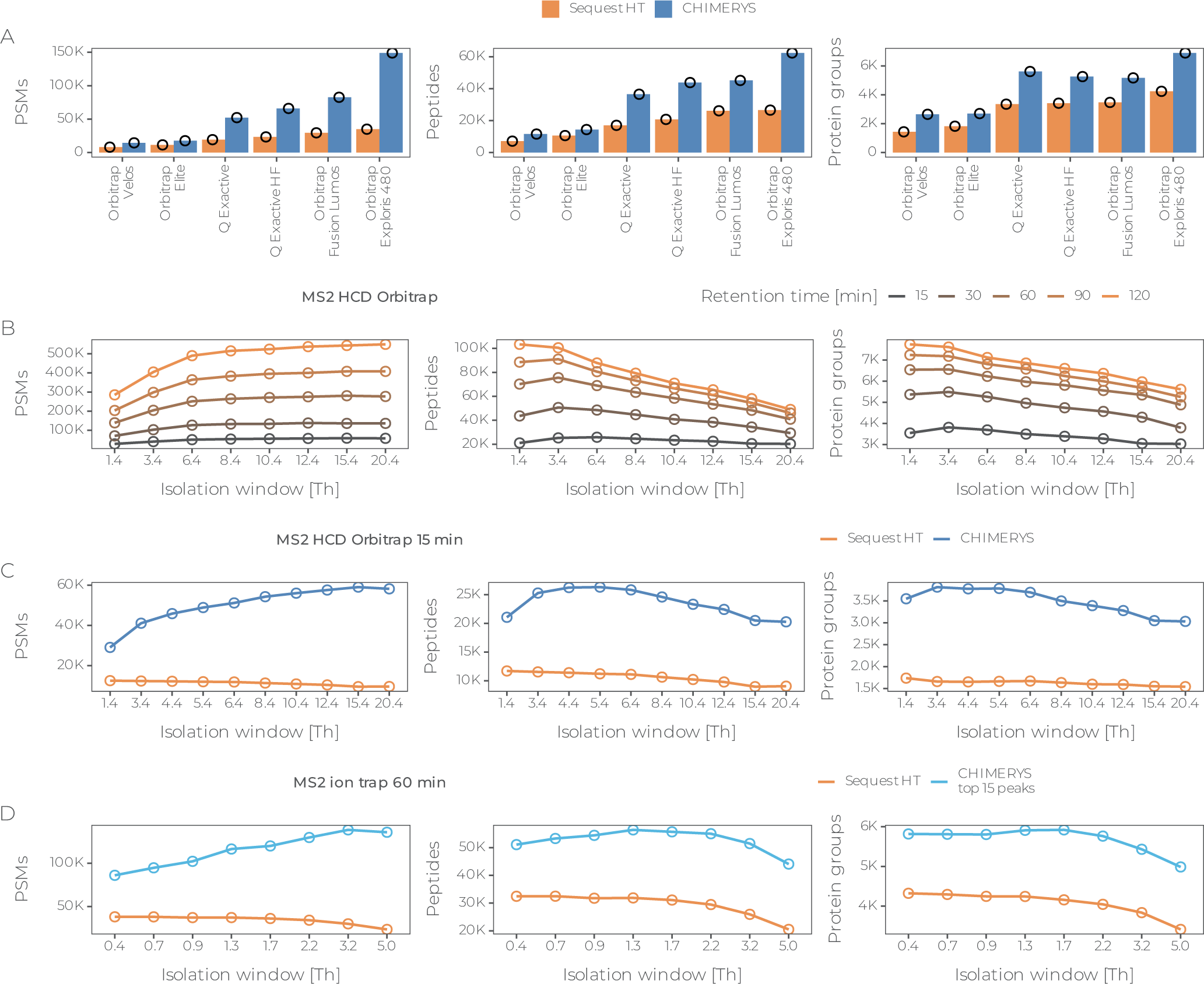

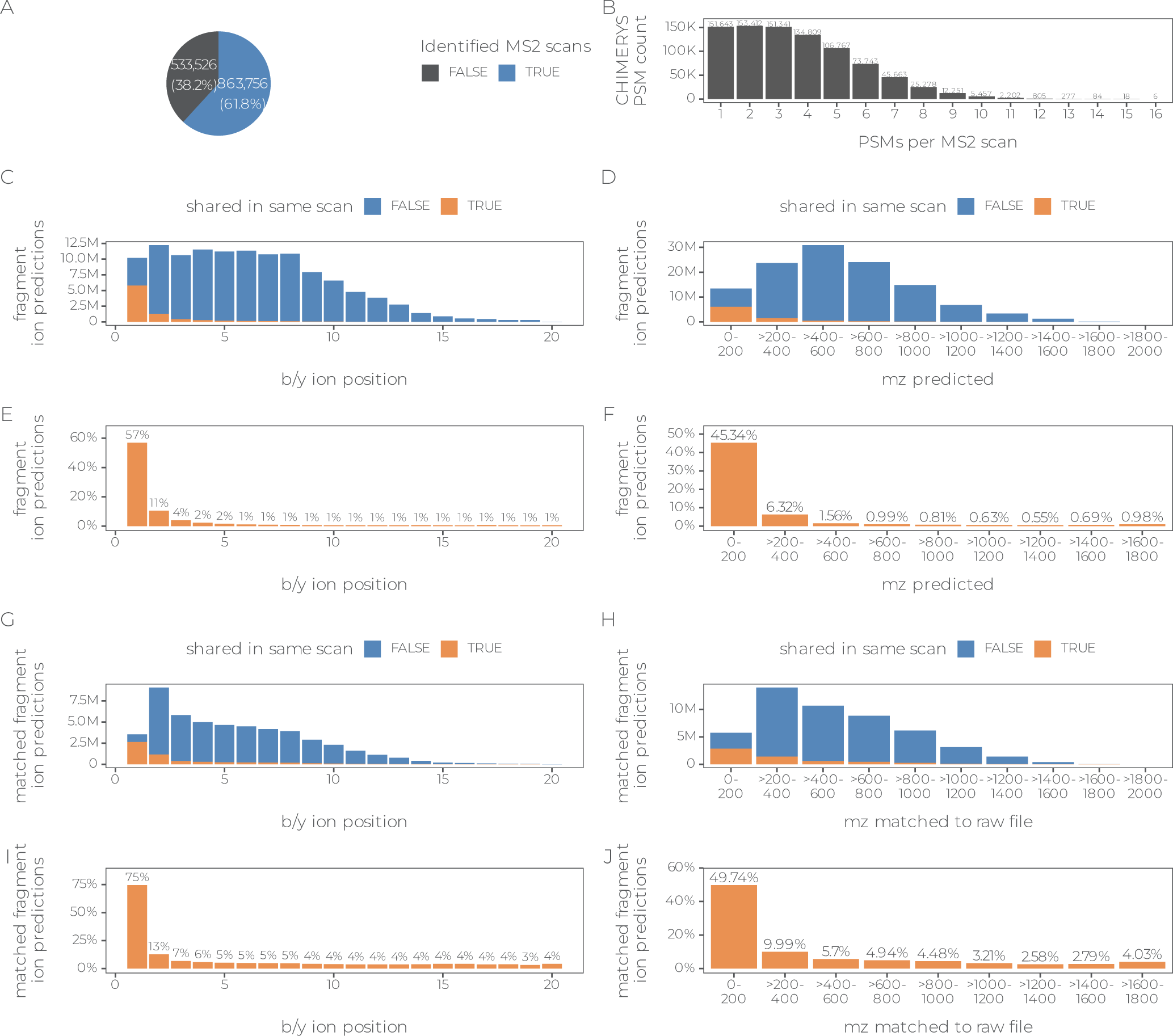

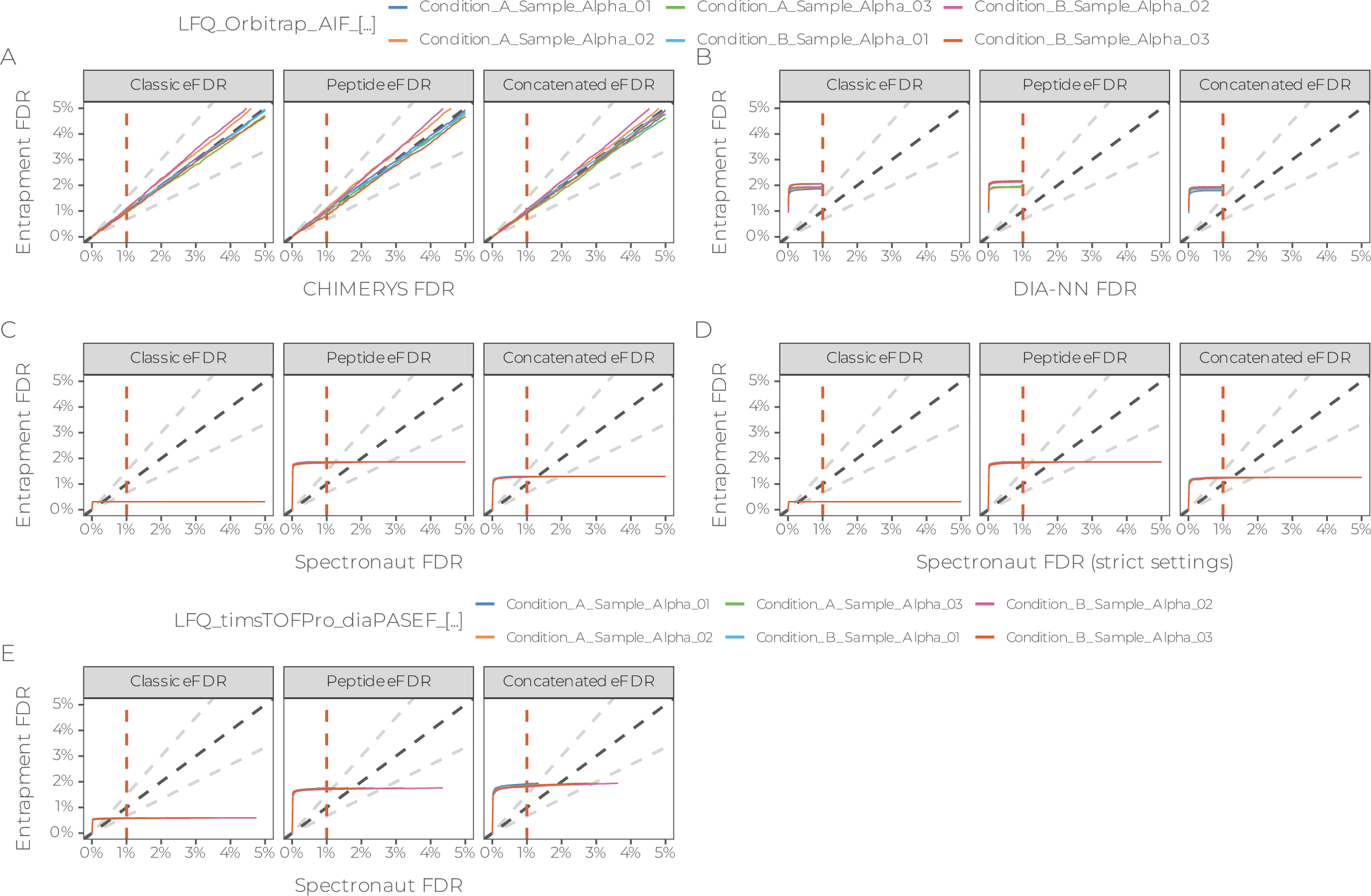

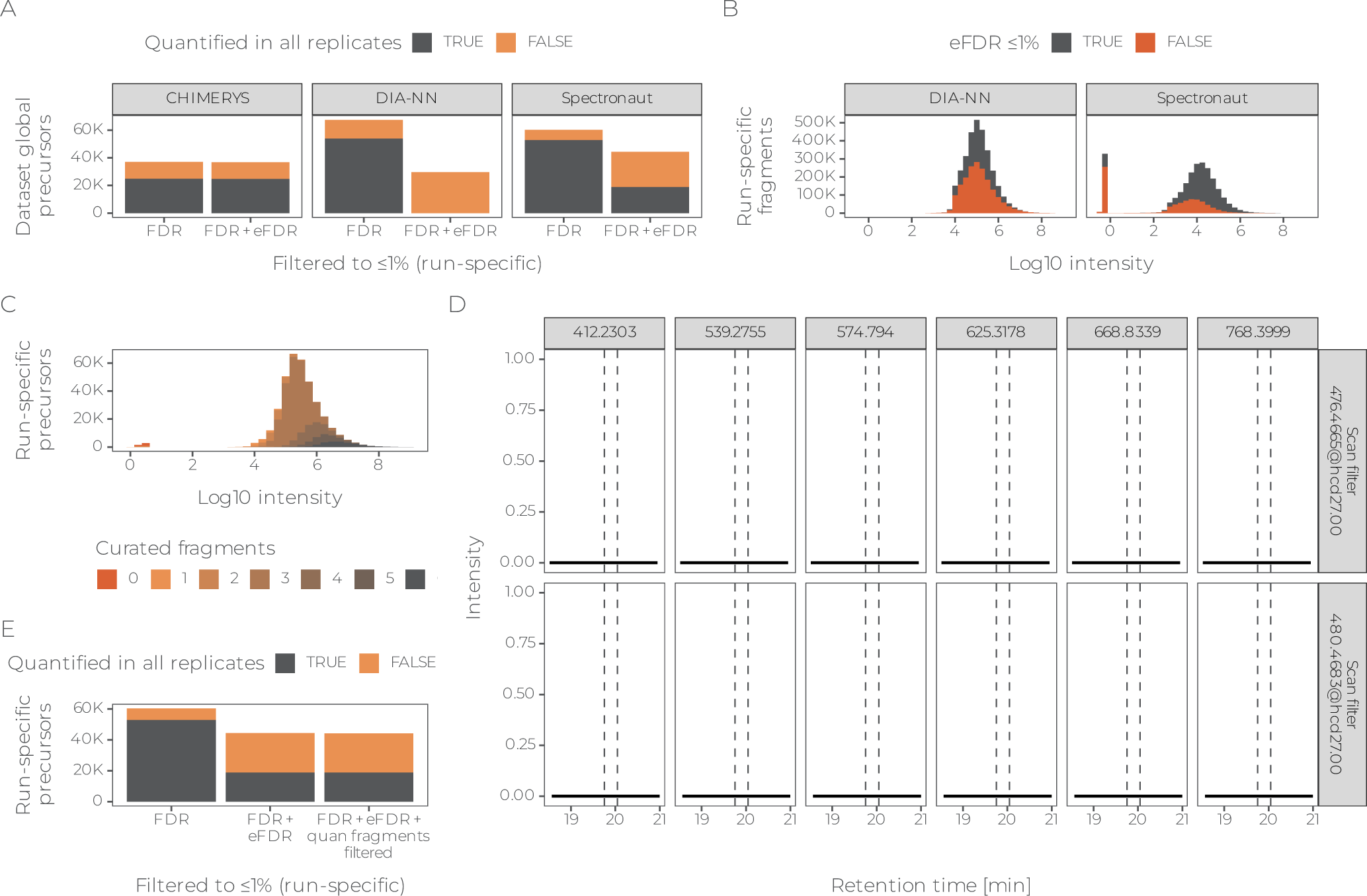

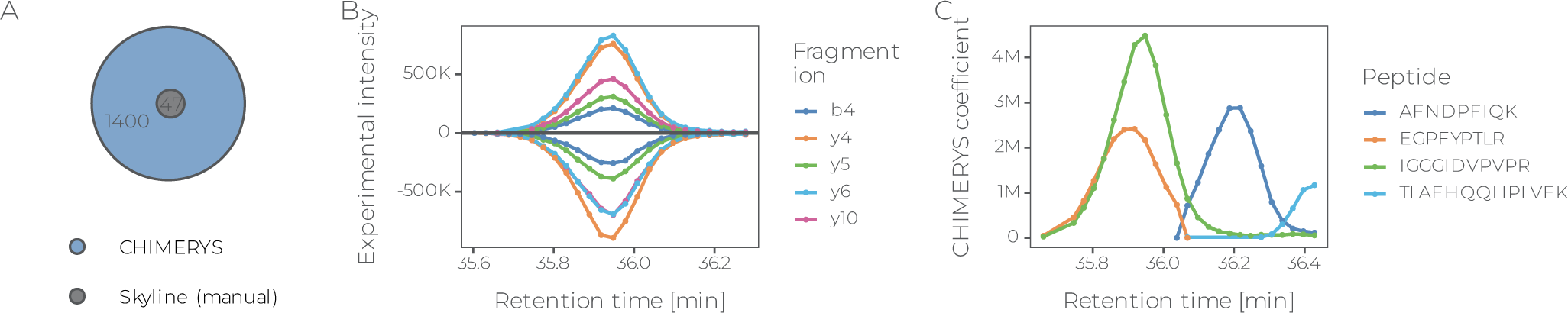

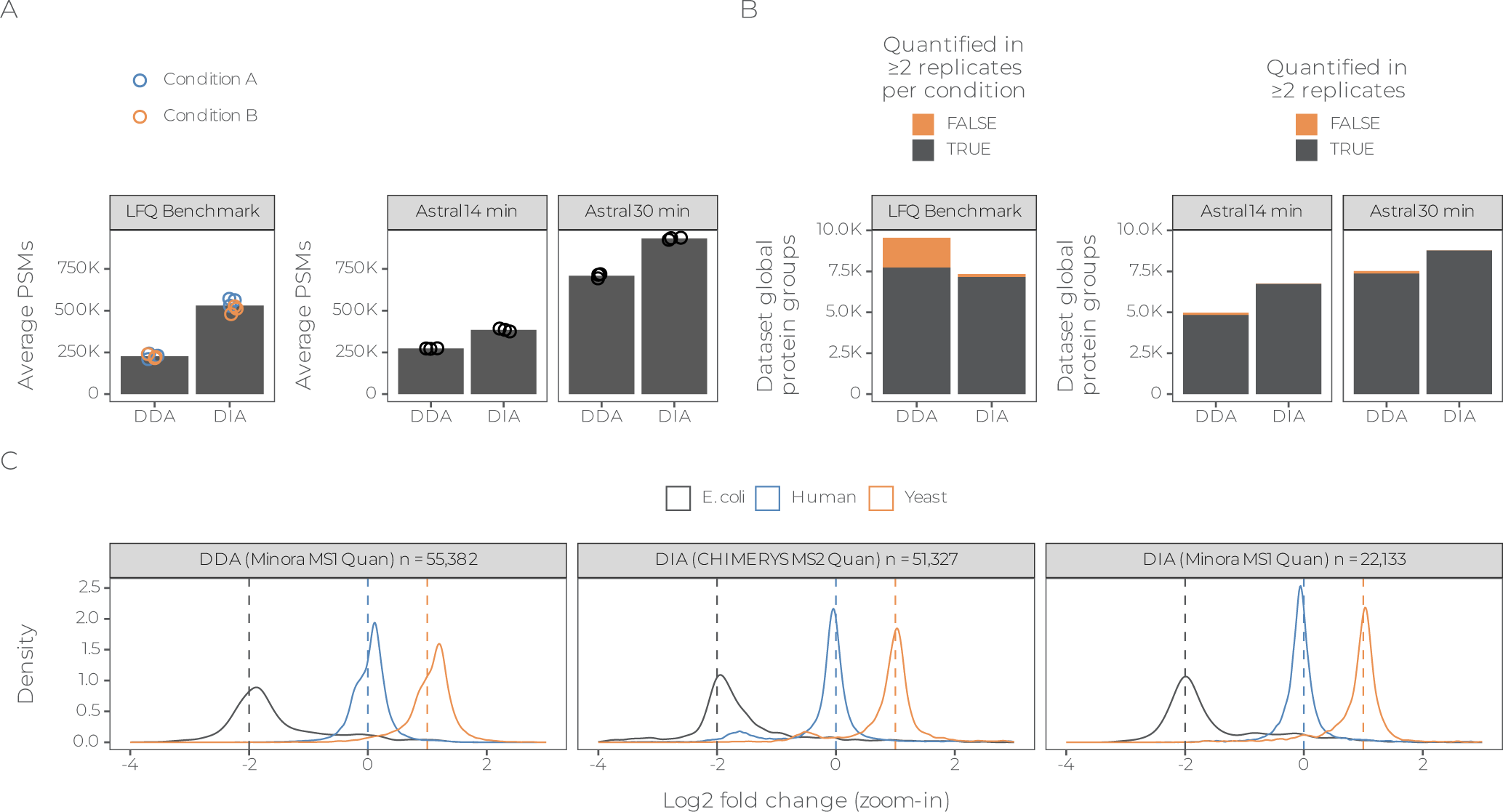

