## Supplementary Information for "Unifying the analysis of bottom-up proteomics data with CHIMERYS"

#### Supplementary Discussion

We have demonstrated that our novel spectrum-centric search algorithm CHIMERYS is capable of deconvoluting chimeric spectra with accurate and sensitive PSM-level FDR control, irrespective of isolation window width, making it suitable for the analysis of DDA, DIA and PRM data alike. CHIMERYS is mindful of the fact that MS2 spectra can contain fragment ions from peptides the theoretical m/z of which falls into a corresponding isolation window, even if the precursor ion was not detected in the preceding MS1 spectrum. This frequently happens due to the limited dynamic range of MS1 spectra that acquire data across a large mass range and thus quickly reach the maximum number of ions that can be captured. MSFragger-DIA[1] takes a similar approach, however, its greedy algorithm that prevents experimental fragment ion peaks to contribute to the score of multiple different peptides is a subtraction approach, which limits its sensitivity, especially close to the detection limit, where typically only few fragment ions are matched, and some of which might be shared (e. g. b2- and y1-ions).

CHIMERYS' novel deconvolution-based approach allows it to distribute the experimental intensity of shared fragment ions to multiple peptides, proportional to their predicted contribution to the total ion current of the experimental spectrum. As a result, CHIMERYS maximizes the amount of explained experimental intensity with a minimal set of peptides. Merely ignoring low m/z regions to avoid shared fragment ions – as often done by DIA search engines – is not sufficient to protect against calling multiple identifications on the same signal. Additionally, CHIMERYS only reports the best scoring PSM per theoretical precursor mass for a given MS2 spectrum to further limit the number of highly similar (and isobaric) peptides being identified based on the same fragment ion information. In a way, this approach returns to the roots of bottom-up proteomics, since CHIMERYS emphasizes the value of PSMs as the central information currency. Its deconvoluted spectra are tangible for the researcher as PSMs can be visually inspected using mirror plots. As it stands, CHIMERYS' acquisition method-agnostic data analysis approach requires highly accurate fragment ion intensities as obtained from pure peptide MS2 spectra, which is why predicted spectra are preferred over experimental spectral libraries, which themselves can contain chimeric spectra. Additionally, using predicted spectra as priors for peptide identification has the advantage that both target and decoy spectra can be predicted with equal quality, ensuring fair competition between targets and decoys. However, the requirement for predicted MS2 spectra is also a limitation of CHIMERYS, since it prevents its application to peptides carrying modifications that cannot yet be predicted by our deep learning model INFERYS. In the future, this limitation should be lifted as deep learning models start to emerge that are capable of generalizing to previously unseen modifications[2].

As part of its deconvolution, CHIMERYS reports coefficients, which are spectrum-centric measures of peptide quantity and can effectively be interpreted as the interference-corrected total ion current of a given peptide. Other approaches typically select a fixed number of the most abundant fragment ions that do not suffer from interference (i. e. do not occur in more than one peptide) for quantification. This has the disadvantage that fragment ion interference can be highly sample specific, potentially resulting in different fragment ions being used for the quantification of the same peptide in different samples. Worst case, this can lead to wrong biological conclusions, especially when analyzing heterogeneous sample cohorts. Conversely, CHIMERYS' quantification considers all theoretical fragment ions that were used for identification also for quantification. We demonstrated that this novel quantification concept is precise and accurate, closely matches quantification via Skyline on PRM data and is highly correlated to quantification in MS1. As such, CHIMERYS is well-equipped for the analysis of heterogeneous sample cohorts such as population-scale plasma or cancer proteomics, as well as single-cell proteomics.

Nowadays, data on such cohorts is often acquired using DIA, since it promises deep proteome coverage, high reproducibility and – most importantly – high data completeness given its systematic way of acquiring MS2 spectra. We have shown that entrapment experiments are important in order to avoid overestimating data completeness exemplified by DIA data analyzed with DIA-NN or Spectronaut. It is worth noting that we do not claim that all precursors not surviving eFDR thresholds are wrong identifications, but rather that our entrapment experiments suggest that neither DIA-NN nor Spectronaut can claim them as confidently identified at 1% precursor-level FDR in the run-specific context.

Notably, Spectronaut quantified some precursors with low self-reported q-values but high eFDR and intensities close to zero. Inspecting fragment-level intensities of precursors identified by Spectronaut revealed that some fragment ions showed peak areas below 1 (Supplementary Figure 9B) and precursors with intensities close to 0 were quantified exclusively based on such fragment ions (Supplementary Figure 9C). Inspection of extracted ion chromatograms for such precursors – an example of which is shown in Figure 3D (bottom) – showed that they were quantified based on fragments with intensities between 0 and 1 across the putative elution of the precursor, which is in stark contrast to similarly high-scoring identifications shared between the three different search engines (Figure 3D top). Inspection of the corresponding raw data showed that these fragment ions sometimes had no signal at all at the relelvant retention time even though Spectronaut reported six data points per peak for the corresponding precursor, implying that Spectronaut performs some signal imputation even when imputation is explicitly turned off (Supplementary Figure 9D). We would argue that such fragment ions should be filtered out and also to remove precursors with fewer than three fragment ions used for quantification as initially configured in Spectronaut (Online Methods).

Interestingly, Spectronaut behaved markedly differently when analyzed with the classic eFDR approach compared to the peptide and concatenated eFDR approaches (see Online Methods). We hypothesize that this is due to the fact that entrapment identifications – unlike random false target identifications – are not randomly distributed across proteins using the classic eFDR approach, but rather always map to entrapment proteins, which – unlike proteins from the original target database – only contain false identifications. Therefore, entrapments generated with the classic eFDR approach can be trivially distinguished from random false target identifications by search engines that incorporate protein-level information into the precursor-level scores used for FDR estimation. An example of such a precursor-level score would be the average intensity of precursors that uniquely map to a given protein. Attaching such a score to each of the precursors mapping to the same protein will boost the confidence in true (non-random) and false (random) precursors mapping to true target proteins, because true precursor identifications will be based mostly on actual signal, while false precursor identifications will be based mostly on noise. Consequently, this would allow precursors mapping to true target proteins to score systematically higher than precursors mapping to entrapment proteins. If this happens during the generation of a spectral library, which is usually filtered to 1% FDR, sometimes at multiple levels, then there is a chance of removing entrapment precursors, but not random false target precursors from said library. This discrepancy between random false target precursors and entrapment precursors is rectified with the concatenated eFDR and peptide eFDR approaches, which is desirable when evaluating precursor-level FDR estimates. As such, measurements of empirical precursor-level FDR with the concatenated eFDR and peptide eFDR approaches are more robust measurements of the true precursor-level FDR. DIA-NN seems to be anti-conservative in its FDR estimates, independent of the entrapment approach used. In fact, DIA-NN loses substantially more data completeness at peptide eFDR than CHIMERYS or Spectronaut. One explanation for this could be that the decoy library entries DIA-NN generates may be too dissimilar from target library entries. Notably, entrapments generated using mimic[3] can be very similar to targets, making it harder to distinguish them from each other, which is a desirable property for entrapments.

While a comprehensive evaluation of all DIA search algorithms is beyond the scope of this paper, we hypothesize that failure to control run-specific precursor-level FDR is due to the way spectral libraries are constructed in library-free workflows. Generating an FDR-filtered spectral library on DIA data and subsequently using it for the peptide-centric scoring of the same data could be considered double dipping, since information on the separation between targets and decoys is incorporated into the spectral library generation, which will in turn influence the peptide-centric FDR estimation performed later. As a result, the q-values of true and false peptide identifications could get boosted to a point where it is actually no longer possible to distinguish between true and false identifications. This would be particularly problematic if samples are highly heterogeneous. The more distinct the peptide set detectable in each sample, the more false identifications could potentially be introduced. Unfortunately, the widespread assumption that DIA measurements have high data completeness due to the way the data was acquired prompts researchers to expect very high data completeness (say 95% or higher), even when analyzing single cells, which are very heterogeneous almost by definition. This issue is exacerbated by novel mass spectrometers that drive sensitivity to the point of single ion detection. Recently, researchers began reporting record numbers of identifications on single cells with almost full data completeness. However, such data completeness claims need to be backed by strong evidence, otherwise, biological conclusions of individual experiments and the reputation of the entire DIA community are at risk.

In a maturing field of proteomics, where soon more biologists than MS experts utilize the technology for advancing research without inspecting the underlying raw data, software developers need to ensure that the output of their tools can be trusted and used at face value. This also creates the necessity to establish reporting guidelines for journals, minimal requirements for calling identifications (e.g. minimum number of matched fragment ions), standardized benchmark measures for evaluating search algorithms and clear community standards for validating error control in DIA and other bottom-up proteomics data. Molecular and Cellular Proteomics has published such guidelines (https://www.mcponline.org/dia-guidelines), but these need to be overhauled to prevent a data reproducibility crisis in proteomics. The new entrapment approaches proposed herein are one contribution to managing quality and several more can be envisaged. Applying such tools will help software developers to meet the responsibility entrusted to them by their users.

In summary, we demonstrated that CHIMERYS properly controls the run-specific precursor-level FDR, while its deconvolution approach makes it agnostic to the chosen data acquisition method, which very timely ties into recent hardware developments, where smaller and smaller DIA windows are closing in on isolation widths typically used for DDA data acquisition. This calls for a harmonization of data analysis strategies, as the differences between DDA and DIA begin to go away[4]. The algorithm presented herein is well-equipped to be able to deal with any current and new acquisition methods and hence unifies the processing of bottom-up proteomics data.

### Supplementary Figure Legends

#### Supplementary Figure 1 – Chimeric DDA spectra

**(A)** Proportions of MS2 spectra with at least one (blue) or no PSM identification (gray) in a 2-hour HeLa DDA single-shot measurement, acquired on an Orbitrap QE HF-X with 1.3 Th isolation windows from the LFQbench-type dataset[5]. **(B)** Distribution of the number of PSMs per MS2 spectrum for the same data as in **(A)**.

#### Supplementary Figure 2 – Shared fragment ions

Absolute number **(A-B)** and relative fraction **(C-D)** of shared (orange) and unshared (blue) fragment ions between predicted spectra of PSMs identified by CHIMERYS at 1% run-specific PSM FDR in a 2-hour HeLa DDA single-shot measurement, acquired on an Orbitrap QE HF-X with 1.3 Th isolation windows from the LFQbench-type dataset as a function of fragment ion position **(A&C)** and fragment ion m/z **(B&D)**. **(E-F)** Analogous plots as in **(A-D)**, but for shared (orange) and unshared (blue) fragment ions from predicted spectra that were matched to experimental peaks.

#### Supplementary Figure 3 – Overlap in identifications

Venn diagram of PSM **(A)** and peptide group **(B-C)** identifications in a 2-hour HeLa DDA single-shot measurement, acquired on an Orbitrap QE HF-X with 1.3 Th isolation windows from the LFQbench-type dataset comparing CHIMERYS to the combination of other search engines at 1% PSM-level FDR **(A)**, 1% peptide group-level FDR **(B)** or 1% FDR at different levels **(C)**. The different FDR levels in **(C)** were the peptide group level for CHIMERYS, Sequest HT, Comet, MS Amanda and MaxQuant, the precursor level for MSFragger and the PSM level for Metamorpheus and MS-GF+.

#### Supplementary Figure 4 – CHIMERYS-unique identifications and FDR levels

**(A)** MS1 apex intensity distribution of peptide groups identified uniquely by CHIMERYS (blue) or also by at least one other search engine tested as part of Figure 1E (gray) at 1% global peptide group-level FDR in a 2-hour HeLa DDA single-shot measurement, acquired on an Orbitrap QE HF-X with 1.3 Th isolation windows from the LFQbench-type dataset. **(B)** Distribution of the number of matched peaks between predicted and experimental spectra for the same data as in **(A)**. **(C)** Distribution of the normalized spectral contrast angle after deconvolution between predicted and experimental spectra for the same data as in **(A)**. **(D)** Scatter plot of the number of unique peptides per protein group identified by SequestHT (x-axis) or CHIMERYS (y-axis) for the same data as in **(A)**. Protein groups are filtered to 1% global protein FDR and peptides are filtered to 1% global peptide group-level FDR. **(E)** The number of precursors (left) or peptide groups (right) identified by CHIMERYS at 1% run-specific PSM- and precursor-level FDR or global peptide group-level FDR in a single 2-hour HeLa DDA single-shot measurement, acquired on an Orbitrap QE HF-X with 1.3 Th isolation windows from the LFQbench-type dataset. **(F)** Same as **(E)**, but for 2-hour DDA single-shot measurement from two different conditions, acquired in three replicates on an Orbitrap QE HF-X with 1.3 Th isolation windows from the LFQbench-type dataset.

#### Supplementary Figure 5 – CHIMERYS-unique identifications and FDR levels

PSM, peptide and protein group identifications based on Sequest HT (orange) and CHIMERYS (blue) from measurements of human urine **(A)**, CSF **(B)** and plasma[6] **(C)**, FFPE **(D)** and secretome samples[7]**(E)**, as well as from publicly available 1h measurements of *Arabidopsis thaliana* and *Halobacterium*[8] **(F)** FDR was controlled at the run-specific PSM-, global peptide group- and global protein level, respectively.

#### Supplementary Figure 6 – Instrument generations, gradients and wwDDA/WWA

**(A)** PSM, peptide and protein group identifications based on Sequest HT (orange) and CHIMERYS (blue) from 1-hour HeLa single-shot measurements, acquired using various Orbitrap generations. FDR was controlled at 1% at the run-specific PSM-, global peptide group- and global protein level, respectively. **(B)** PSM, peptide and protein group identifications based on CHIMERYS from HeLa single-shot measurements, acquired using different gradient lengths and isolation window widths. FDR was controlled at 1% at the run-specific PSM-, global peptide group- and global protein level, respectively. **(C)** PSM, peptide and protein group identifications based on Sequest HT (orange) and CHIMERYS (blue) from 15 min HeLa single-shot measurements, acquired using HCD fragmentation with Orbitrap readout and different isolation window widths. FDR was controlled at 1% at the run-specific PSM-, global peptide group- and global protein level, respectively. **(D)** PSM, peptide and protein group identifications based on Sequest HT (orange) and CHIMERYS after removal of low-abundance peaks (light blue) from 1-hour HeLa single-shot measurements, acquired using CID fragmentation with ion trap readout and different isolation window widths. FDR was controlled at 1% at the run-specific PSM-, global peptide group- and global protein level, respectively.

#### Supplementary Figure 7 – Chimeric DIA spectra

**(A)** Proportions of MS2 spectra with at least one (blue) or no PSM identification (gray) in triplicate 2-hour DIA single-shot measurements from two different conditions, acquired on an Orbitrap QE HF-X with 8 Th isolation windows from the LFQbench-type dataset[5]. **(B)**

Distribution of the number of PSMs per MS2 spectrum for the same data as in **(A)**. **(C-D)** Absolute number and relative fraction **(E-F)** of shared (orange) and unshared (blue) fragment ions between predicted spectra of PSMs identified by CHIMERYS at 1% run-specific PSM FDR in the same data as in **(A)** as a function of fragment ion position **(C&E)** and fragment ion m/z **(D&F)**. **(G-J)** Analogous plots as in **(C-F)**, but for shared (orange) and unshared (blue) fragment ions from predicted spectra that were matched to experimental peaks.

#### Supplementary Figure 8 – Entrapment analyses on DIA data

Scatter plots of run-specific precursor-level self-reported (x-axis) and entrapment FDR (y-axis) from three different entrapment approaches (Online Methods) for CHIMERYS **(A)**, DIA-NN **(B)**, Spectronaut with default settings **(C)** and Spectronaut with more stringent settings[9] **(D)** on triplicate 2-hour DIA single-shot measurements from two different conditions, acquired on an Orbitrap QE HF-X with 8 Th isolation windows from the LFQbench-type dataset[5]. **(E)** same as in **(C)**, but for the corresponding TimsTOF Pro data.

#### Supplementary Figure 9 – DIA data analysis with CHIMERYS, DIA-NN and Spectronaut

**(A)** Precursors quantified by CHIMERYS, DIA-NN and Spectronaut in at least one (orange) or three (gray) out of three replicate measurements of two different conditions in a multispecies LFQbench dataset. Identifications are filtered at 1% run-specific precursor-level FDR or additionally also at 1% run-specific precursor-level entrapment FDR based on the peptide eFDR approach (Online Methods). **(B)** Peak areas (F.PeakArea) for fragment ions from precursors surviving (gray) or not surviving (red) 1% run-specific precursor-level entrapment FDR based on the peptide eFDR approach for the same data as in **(A)**, analyzed by DIA-NN and Spectronaut. **(C)** Apex intensities for precursors identified by Spectronaut at 1% run-specific precursor-level FDR for the same data as in **(A)**, colored by the number of fragment ions with Peak areas (F.PeakArea) > 1 that were not excluded from quantification by Spectronaut. **(D)** Example fragment ion XICs directly extracted from the raw file for the precursor at m/z 479.2478 identified by Spectronaut but not by CHIMERYS in Figure 3D (all six library fragments are shown). XICs were extracted using the R package rawrr with a fragment mass accuracy of 20 ppm. **(E)** Precursors quantified by Spectronaut in at least one (orange) or three (gray) out of three replicate measurements of two different conditions in a multispecies LFQbench dataset. Identifications are filtered at 1% run-specific precursor-level FDR, additionally also at 1% run-specific precursor-level entrapment FDR based on the peptide eFDR approach (Online Methods) and additionally also by excluding precursors that are quantified based on less than three fragment ions with peak areas (F.PeakArea) > 1, which were not excluded from quantification by Spectronaut.

#### Supplementary Figure 10 – PRM and DIA quantification using CHIMERYS coefficients

**(A)** Venn diagram of peptides identified in a PRM dataset – targeting 52 peptides from 18 human proteins – by CHIMERYS at 1% precursor-level FDR (blue) or Skyline (gray). **(B)** Mirror XIC of the top five experimental (above the x-axis) and predicted fragment ion intensities, scaled by the corresponding CHIMERYS coefficients (below the x-axis) for one of the targeted peptides in A. **(C)** Coefficient-based reconstruction of elution peaks for four different peptides identified by CHIMERYS in the data in **(A)**, only one of which was targeted in the assay (IGGGIDVPVPR).

#### Supplementary Figure 11 – Comparisons of DDA and DIA data

Precursors **(A)** and protein groups **(B)** identified by CHIMERYS in triplicate 2h single-shot measurements from a multispecies LFQbench dataset acquired in DDA (blue) or DIA (orange) on an Orbitrap QE HF-X (left) or in triplicate 30min single-shot measurements from a HeLa sample acquired in DDA (blue) or DIA (orange) on an Orbitrap Astral (right). FDR was controlled at 1% at the run-specific PSM-level or at the global protein level, respectively. Match between runs was used for DDA data and for DIA data, peptides were quantified irrespective of their run-specific FDR. **(C)** Peptide-level $\log_2$-ratio density plots for the same DDA and DIA data from the LFQbench dataset as in **(A)**, quantified in MS1 using the Minora Feature Detector and in MS2 using CHIMERYS.
